# Introducing a fusogenicity metric for lipid nanoparticle formulation

**DOI:** 10.64898/2026.03.02.708638

**Authors:** Lining Zheng, Max Baliga, Seamus F. Gallagher, Alice Gao, Jacob Rueben, Yoo Kyung Go, Markus Deserno, Cecilia Leal

## Abstract

Lipid nanoparticles (LNPs) are the most successful drug delivery carrier to date, but optimizing lipid formulations to improve membrane fusion capabilities for effective drug release has been challenging due to lack of a quantitative measure for fusogenicity. Here we introduce a new framework based on small angle X-ray scattering to experimentally measure 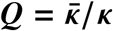 for lipids used in LNP formulations such as glycerol monooleate (GMO) and ionizable lipids (SM-102 and ALC-0315). *Q* intrinsically captures spontaneous curvature (*J*_0_), which is traditionally used to assess fusogenicity. The change of cubic lattice parameters with temperature was measured for GMO-containing lipid mixtures, and the *Q* extracted quantitatively correlated with LNP fusogenicity power validated by fluorescence-based fusion assays and cryogenic electron microscopy. Fusogenicity of SM-102 and ALC-0315 was quantified by adding them to host membranes and assessing change in *Q*. This framework provides researchers with the ability to optimize the fusogenicity of LNP formulations for potent drug release and enhances understanding of parameters governing fusion in all biomembranes.

---

Lipid nanoparticles (LNPs) are the most successful nanocarriers to date, demonstrated by their use in the COVID-19 mRNA vaccines that saved an estimated 14.4 million lives (*1*). Despite their success, LNPs still suffer from major limitations, including off-target effects, immunogenicity, and importantly very low delivery efficiency (*2*) (*3*) (*4*). Endocytosis is the preferred mode of cell entry for most viral and non-viral delivery systems (*5*) (*6*) (*7*) (*8*) (*9*). Our current understanding of LNP delivery mechanisms pins low delivery efficiency to LNP entrapment inside endosomal vesicles (*10*) (*11*). As the endosome matures, it enters the lysosomal pathway and the content entrapped is degraded. LNPs need to release their cargo into the cytosol in a timely manner (less than 30 min to 1 hour) (*12*) to prevent the degradation of their payload. Viral proteins have the capacity to change conformation in the endosomal milieu that boosts the rapid formation of a fusion pore between the viral envelope and the endosomal membrane through which cargo is released. Compared to viral delivery vehicles, LNPs require vigilant engineering to effectively escape the endosome and deliver payload. Enhancing the ability of LNPs to fuse with endosomes, i.e. increasing LNP *fusogenicity*, and forming small fusion pores is the most promising way to boost delivery efficiency while minimizing disruption of intracellular membranes that may lead to uncontrolled cellular responses, such as increased inflammation (*13*). However, methods to predict and quantify LNP *fusogenicity* remain critically elusive. We do not have a robust methodology to predict fusogenicity of novel ionizable lipids with promising therapeutic potential (*14*) (*15*) (*16*) (*17*) (*18*) (*19*) (*20*) (*21*). Fusogenicity assays based on fluorescence resonance energy transfer (FRET) to track LNP fusion with model endosomal membranes have been enabling (*22*) (*23*) (*24*), but only provide a relative comparison of fusogenicity between formulations, not a quantitative measure. Moreover, these experiments require extensive experimental optimization for parameters such as the fluorescent dye concentration and ratio. Attempting to explore the pH-dependent behavior of ionizable lipids further complicates the issue.

In this paper, we develop the first quantitative framework to robustly predict the fusogenicity power of any lipid synthesized for LNP formulation. More broadly, our method uncovers new physical insights of the fundamental parameters that control membrane fusion, a prevalent process in all biomembranes. Specifically, we highlight that the elastic energy cost of lipid membrane fusion and fusion pore formation is strongly dependent on its inherent Gaussian modulus 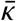. The membrane elastic energy can be represented by the Helfrich equation (*25*) (*26*),

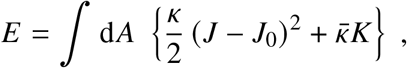

where *κ* is the bending modulus, *J* is the total/extrinsic curvature which equals the sum of the principle curvatures *C*_1_ + *C*_2_, and *J*_0_ is the spontaneous curvature. The second term comprises the Gaussian curvature *K* = *C*_1_*C*_2_ and the Gaussian modulus 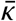. By using the Gauss-Bonnet theorem (*27*), we can deduce from Helfrich theory that the elastic energy cost of forming a membrane pore 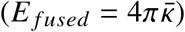 from two separate membranes 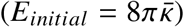 is 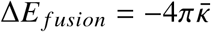 (*28*). As schematically introduced in Fig. 1, it is clear that the Gaussian modulus 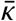 is the main deciding factor on how easily the lipid membrane of an LNP fuses with the endosomal membrane. Unfortunately, while several methods are available to measure *κ* (*29,30*) such as fluctuation analysis or micropipette aspiration, the Gaussian modulus 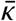 is difficult to measure experimentally and computationally (*31*)(until very recently at least). Due to the mentioned Gauss-Bonnet theorem, the Gaussian modulus would only be relevant when a topological transformation occurs (*32*). These transformation processes are hard to control experimentally, making it difficult to extract 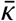. Only few 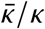 values have been published (*32*). One promising method introduced in decades past relies on varying the lattice parameter of bicontinuous cubic phases to water content (*33*) or temperature (*34*) and creating giant liposomes with fluid phase coexistence (*35, 36*). However, these studies only evaluated the 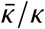 of one or two lipids or lipid mixtures, and did not demonstrate correlation of 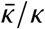 with the fusion behaviors of LNPs with actual endosomal membranes.

**Figure 1:**
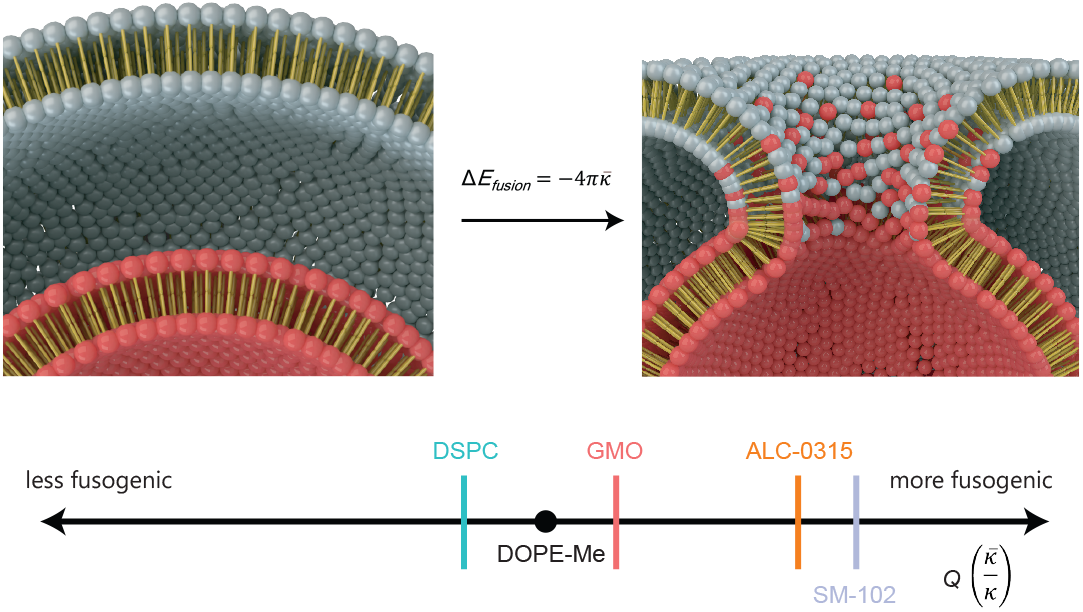
The elastic energy cost of forming a membrane pore depends on 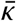, **and** 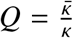 can be compared across different lipids with our platform. Illustrations credits: Alex D. Jerez, Imaging Technology Group at the Beckman Institute University of Illinois, Urbana-Champaign, IL.

We have previously shown that LNPs made of high molar percentages of glycerol monooleate (GMO) can form bicontinuous cubic phase LNP-RNA complexes (cuboplexes) and that cuboplexes are able to escape the endosome more efficiently compared to lamellar LNP-RNA complexes (lipoplexes) (*22*) (*37*) (*38*) (*39*) (*40*) (*41*) (*42*) (*43*) (*44*). However, this work only qualitatively evaluates cuboplex fusogenicity.

In this paper, we define a new fusogenicity parameter *Q*, the Gaussian modulus 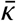 normalized to bending modulus 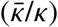 of lipid bilayers, yielded from SAXS measurements of the lattice parameters of bicontinuous phases at different temperatures. Importantly, *Q* intrinsically encodes lipid packing parameter (*45*), via the associated spontaneous curvature *J*_0_, which has been elusively used a predictor of fusogenicity (*46*) (*47*) (*48*) (*49*). The predictive power of *Q* as a *fusogenicity* parameter was validated by FRET-based assays carried out for a series of LNPs at different compositions. In addition, membrane fusion contacts between LNPs and endosomes were evaluated by cryogenic electron microscopy (cryo-EM). Our results show that increasing GMO content can increase the fusogenicity of LNPs. We also found that the addition of ionizable lipids that are used in the COVID-19 vaccines developed by Moderna (SM-102) and Pfizer/BioNTech (ALC-0135) have a significant effect on *Q* of a host membrane. Such effects are enhanced at low pH when the ionizable lipids are protonated.

Our work underpins the importance of 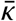 in fundamentally understanding membrane fusion (and fission) processes which are ubiquitous in nature. Specifically, we define *Q* as a robust parameter to predict and quantify fusogenicity power as a critical component in the design of efficient LNP delivery systems.

## Measuring the lattice parameter of *Q*_*II*_ phases vs. Temperature

We start by measuring the fusogenicity parameter *Q* for the special case of lipids that are prone to self-assemble into LNPs having bicontinuous cubic structures *Q*_*II*_ . Next we introduce high contents of cholesterol (30 mol%) that do not disrupt the *Q*_*II*_ phase. Ultimately, we demonstrate that the *Q* metric platform is generalizable to predict *fusogenicity power* of any lipid compound regardless of their ability to form *Q*_*II*_ structures, including ionizable lipids used in clinically relevant LNP formulations.

Our recent work (*22*) (*43*) (*37*) shows that LNPs prepared with high contents of GMO adopt a *Q*_*II*_ structure and are highly fusogenic. The relation between membrane fusion and *Q*_*II*_ phases has been established long ago (*50*) (*51*) (*52*) (*53*) and Siegel and co-workers demonstrated (*34, 54*) that evaluating the shrinkage of the lattice parameter *a* of *Q*_*II*_ phases as a function of temperature yields the monolayer Gaussian modulus, normalized to the bending modulus 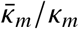. For this study, two sets of GMO formulations were made, one with GMO and DOPC (phosphatidylcholine, 1,2-dioleoyl-sn-glycero-3-phosphocholine) and another including 30% molar percentage of cholesterol (Chol) as high amounts of Chol are used in most LNP formulations (the composition details are shown in Fig. 2a). DPPC (dipalmitoylphosphatidylcholine, 1,2-dipalmitoyl-sn-glycero-3-phosphocholine) and Chol lipid mixtures were also prepared, since they are known to readily form lamellar phases (*55*) (*56*). Samples of these lipid formulations were then made in excess water and flame-sealed in quartz capillaries. The phase and lattice parameter were measured using synchrotron small angle X-ray scattering (SAXS) from 30 °C to 70 °C.

**Figure 2:**
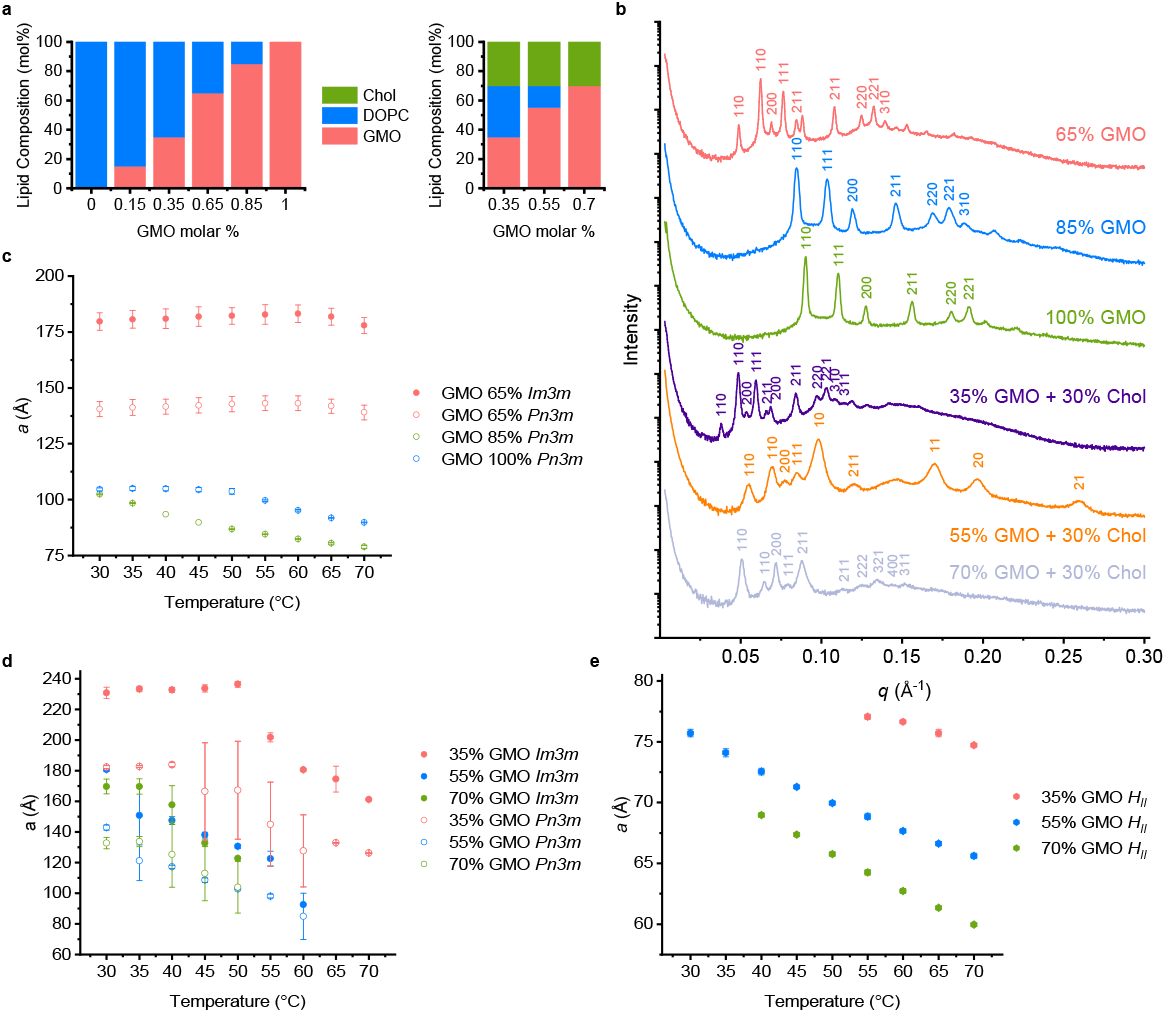
SAXS measurements of GMO/DOPC and GMO/DOPC/Chol mixtures at different temperatures. (**A**) Representation of the composition of the different GMO/DOPC and GMO/DOPC/Chol mixtures used. (**B**) Representative SAXS profiles of the GMO/DOPC and GMO/DOPC/Chol mixtures at 35°C. Miller indices are shown for Bragg reflections of *Q*_*II*_ (*Im*3*m* or *Pn*3*m*) and *H*_*II*_ phases. (**C**) Changes of Lattice parameter with temperature for GMO/DOPC mixtures that have *Q*_*II*_ phases. At higher temperatures, lattice parameter decreases as temperature increases. (**D**,**E**) Changes of Lattice parameter with temperature for GMO/DOPC/Chol mixtures that have *Q*_*II*_ (**D**) and *H*_*II*_ (**E**) phases. Similar to mixtures without Chol, lattice parameter decreases as temperature increases, especially at higher temperatures.

As expected, the DPPC/Chol lipid mixtures formed lamellar phases. As shown in Fig.S1, the lattice parameter decreases with the increase in temperature due to bilayer thinning. When the Chol molar fraction increases, the lattice parameter changes less with temperature change.

For the compositions with GMO/DOPC (Fig.2 A), GMO molar percentages with 0%, 15% and 35% adopt a lamellar phase (*L*) at all the temperatures measured, as shown in Fig.S2. The lattice parameters of these three compositions are similar, ranging from around 63.2 °A to 65.3 °A, and the greater the GMO molar percentage the larger the lattice parameter. For DOPC (GMO 0%), the lattice parameter does not change much with temperature. However, for 15 and 35% GMO the lattice parameter initially increases with temperature due to thermal expansion up to 45 °C, then subsequently decreases. The formulations with GMO molar percentages of 65%, 85% and 100% all display bicontinuous cubic structures (*Q*_*II*_ ) (Fig. 2B,C). GMO molar percentage of 65% adopt both *Im*3*m* and *Pn*3*m* space groups, while GMO molar percentage of 85% and 100% only show the *Pn*3*m* phase. Contrary to the lamellar phase compositions, the greater the GMO molar fraction, the smaller the *Q*_*II*_ lattice parameter.

When 30% cholesterol is included, GMO/DOPC/Chol compositions (Fig. 2B,D) show different behaviors. All the compositions measured showed both inverse hexagonal phase(*H*_*II*_ ) as well as *Q*_*II*_ phases, in which the *Q*_*II*_ phases have both *Im*3*m* and *Pn*3*m* symmetries present. The lattice parameters of *Q*_*II*_ *Im*3*m* phases are larger than those of *Q*_*II*_ *Pn*3*m*. For GMO molar percentage of 35%, the cubic lattice parameters decrease as temperature increases above 50 °C. *H*_*II*_ phases appear starting from 55 °C. For GMO molar percentage of 55%, the cubic lattice parameters decrease as temperature increases above 35 °C. The *H*_*II*_ phase is present at all temperatures tested (Fig. 2E), and from 65 °C the *Q*_*II*_ phase is no longer observed. For GMO molar percentage of 70%, the cubic lattice parameters decrease as temperature increases above 35 °C. For 70% GMO, the temperature at which only the *H*_*II*_ phase is present is lower, at 55 °C. Critically, the *Q*_*II*_ lattice parameters decrease for all samples as the temperature increases.

## Determining 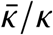 of bilayers from bicontinuous cubic lattice size

The topological transformation of two opposed membranes into a fusion pore is controlled by 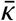 for lipid bilayers. However, while experimental and computational tools to measure bending moduli *κ* are abundant, quantitatively assessing 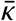 is exceedingly difficult. In simulation, it has been proposed that 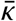 can be related to microscopic transmembrane stresses (*57, 58*). Unfortunately, the values of 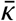 obtained in this fashion disagree with a macroscopic, shape based method that avoids the Gauss-Bonnet theorem by instead simulating open patches (*32, 59*). Experimentally, it is even harder to get around the Gauss-Bonnet theorem. However, the non-trivial topology of cubic phases provides a path forward. Siegel and co-workers (*34*) described an X-ray method where *a*= *f* (*T* ) (see Fig. 2) may be fit to obtain *M*, the ratio of the Gaussian to the bending moduli for monolayers:

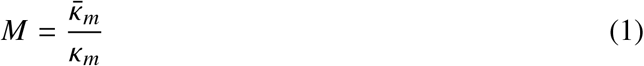

For bicontinuous cubic phases, the unit cell parameter *a* can be expressed as:

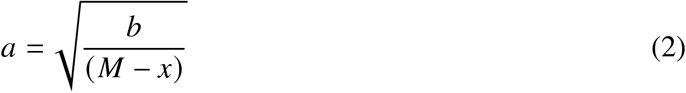

where

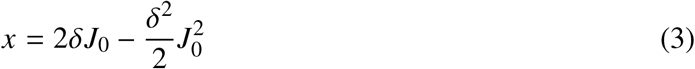

δ is the distance between the bilayer midplanes and the neutral surface of the lipid monolayers. Since both DOPC and GMO are oleoyl-chain lipids, 1.3 nm is used as *δ* throughout this study (*34*) (*60*). The choice of *δ* impacts the *M* value extracted, as is elaborated in subsequent sections. *J*_0_ is the spontaneous curvature of the monolayer, which is temperature dependent. The spontaneous curvature is additive with respect to its molar fraction if the lipids are well-mixed (*61*) (*62*), which is ensured by chloroform solution mixing assuming no phase separation:

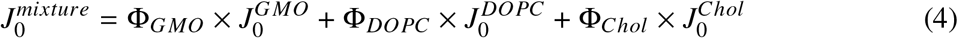

Values of *J*_0_ for DOPC and Chol as a function of temperature are well reported (*63*), but the spontaneous curvature of GMO and its temperature dependence is less available, so we determined experimental values for *J*_0_ using previously established methods (*61*) (*64*) (*63*) (*65*). Since GMO does not spontaneously form monolayers at most temperatures, different amounts of GMO were added to DOPE (1,2-dioleoyl-sn-glycero-3-phosphoethanolamine), which readily forms *H*_*II*_ phases consisting of monolayer tubes hexagonally packed in 2D. Here, the lattice spacing directly relates to the curvature of the monolayer. 9-cis-tricosene was added to relieve packing frustration and relax the *H*_*II*_ phase to form cylindrical tubes. The lattice parameter of the *H*_*II*_ phase was measured by SAXS. By using the pivotal plane of DOPE from previous published values (*65*) (*63*), *J*_0_ of GMO can be extrapolated from the *J*_0_ of different DOPE/GMO mixtures (Fig. S3). By repeating the *J*_0_ measurement for different temperatures, the *J*_0_ of GMO with respect to temperature has been obtained (Fig. S4), shown in Table 1. The *J*_0_ of GMO at 27 °C is calculated to be -0.524 ± 0.01 nm^−1^, which is similar to the -0.54 ± 0.03 at 27 °C published before (*61*). The spontaneous curvatures used for this study are listed in Table 1.

**Table 1:** The spontaneous curvature (*J*_0_) in respect to temperature used in this study. The DOPC and Chol *J*_0_(*T* ) are from reference (*63*) and the GMO *J*_0_(*T* ) was calculated from SAXS measurements of DOPE/GMO mixtures. The temperature (T) is in Celsius (°C).

| Lipid | $J_0(T)$ ($\text{nm}^{-1}$) |
| --- | --- |
| DOPC | $-1.1 \times (T - 35)/1000 - 0.091$ |
| Chol | $-3.5 \times (T - 35)/1000 - 0.494$ |
| GMO | $-0.7 \times T/1000 - 0.505$ |

From the known *M*, we can calculate the ratio of the Gaussian to bending moduli for bilayers 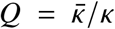. When the total membrane tension is set to zero, 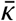 can be expressed as Equation 5 (Derivation shown in Supplementary Text). By dividing both sides of the equation with *κ*_*m*_, and since *κ* is *κ* = 2*κ*_*m*_ (*66*) (*67*), we can derive an expression for *Q* as shown in Equation 6. It is evident that *Q* which we describe in this paper as the *fusogenicity parameter* depends on *M, J*_0_ and *δ*.

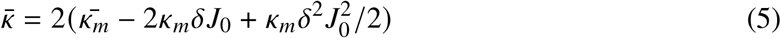

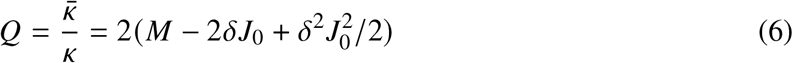

For the GMO/DOPC and GMO/DOPC/Chol compositions that adopt a bicontinuous cubic structure, the *x* values were calculated and the lattice parameter *a* was plotted versus *x*. The data were then fitted for *M* using Equation 2 as shown in Fig. S8,S9. To present the data in a more intuitive manner, Equation 2 can be rearranged into Equation 7. We find that 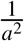 linearly decays with *x* and the intercept *x* (0) is *M*, the ratio of the elastic moduli for monolayers. The 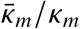 values obtained are shown in Table 2 and Table 3. The fitted functions are shown in Fig. 3A,B for GMO/DOPC and GMO/DOPC/Chol, respectively. It is clear that increasing GMO molar percentage has a strong effect on *M* that becomes more negative (Fig. 3C). It is noteworthy that even with different bicontinuous cubic phases symmetry groups such as *Im*3*m* and *Pn*3*m*, the *M* values are similar for the same lipid composition,, indicating the robustness of this method.

**Table 2:** The 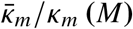 values extracted and *Q* calculated for GMO/DOPC compositions that have cubic phases at 37°C.

| GMO/DOPC Molar Ratio | Phase | $M$ | $Q$ |
| --- | --- | --- | --- |
| 65/35 | $Im3m$ | $-0.719 \pm 0.294$ | $0.383 \pm 0.268$ |
| 65/35 | $Pn3m$ | $-0.711 \pm 0.146$ | $0.392 \pm 0.120$ |
| 85/15 | $Pn3m$ | $-1.277 \pm 0.008$ | $0.116 \pm 0.012$ |
| 100/0 | $Pn3m$ | $-1.462 \pm 0.005$ | $0.156 \pm 0.016$ |

**Table 3:**
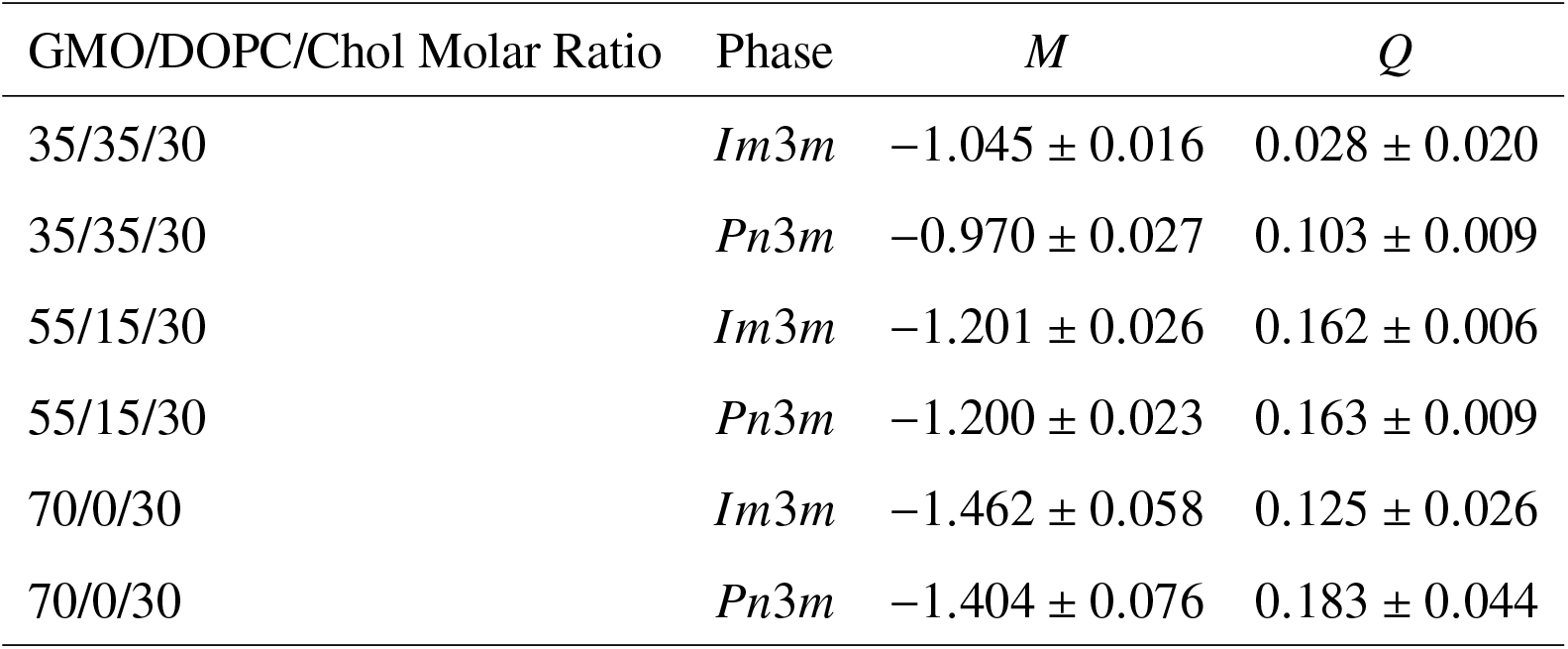
The 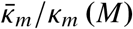 values extracted and *Q* calculated for GMO/DOPC/Chol compositions that have cubic phases.

| GMO/DOPC/Chol Molar Ratio | Phase | $M$ | $Q$ |
| --- | --- | --- | --- |
| 35/35/30 | $Im3m$ | $-1.045 \pm 0.016$ | $0.028 \pm 0.020$ |
| 35/35/30 | $Pn3m$ | $-0.970 \pm 0.027$ | $0.103 \pm 0.009$ |
| 55/15/30 | $Im3m$ | $-1.201 \pm 0.026$ | $0.162 \pm 0.006$ |
| 55/15/30 | $Pn3m$ | $-1.200 \pm 0.023$ | $0.163 \pm 0.009$ |
| 70/0/30 | $Im3m$ | $-1.462 \pm 0.058$ | $0.125 \pm 0.026$ |
| 70/0/30 | $Pn3m$ | $-1.404 \pm 0.076$ | $0.183 \pm 0.044$ |

**Figure 3:**
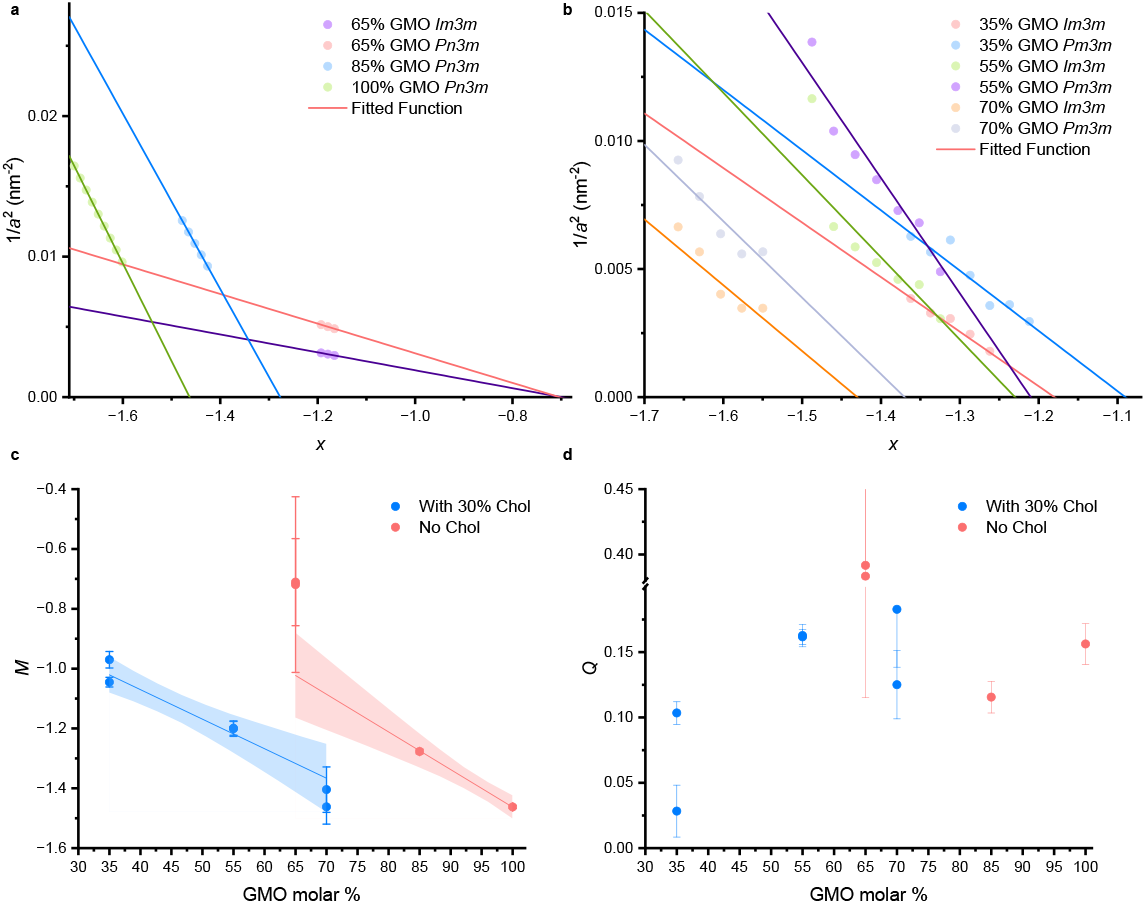
Obtaining the fusogenicity parameter *Q* for GMO/DOPC and GMO/DOPC/Chol mixtures. Plots of 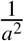 against *x* for GMO/DOPC compositions (**A**) and GMO/DOPC/Chol compositions (**B**), with the fitted linear function where the x intercept is *M*. (**C**) *M* values plotted against GMO molar percentages. *M* becomes more negative as GMO content increases. The red or blue line is a linear regression fit with the 95% confidence interval shown (red or blue shade).(**D**) Fusogenicity parameter *Q*, which is 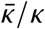, plotted against GMO molar percentages.

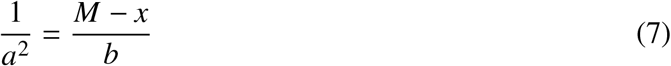

It is tempting to think that the trend is reverse to what is expected as the Gaussian modulus should become less negative for more fusogenic systems at higher GMO content. However, what controls the topological transition associated with the formation of a membrane fusion pore is 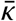 (Gaussian modulus for bilayers). Using equation 6 we calculate *Q* (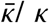 for bilayers) for these compositions and find that indeed *Q* increases with increasing amounts of GMO (Fig. 3 d and Table 2). The elastic energy cost to create a fusion pore is 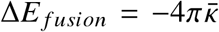 and an increase in *Q* indicates that GMO makes the lipid mixtures more fusogenic. We can also see that at high molar percentages of GMO, *Q* stabilizes and does not increase as significantly, which suggests that fusogenicity plateaus at high molar percentages of GMO.

It is important to note that the choice *δ* significantly influences the *M* values extracted (Fig. S5). In this work the *δ* used is 1.3 nm, but this may not be accurate especially for mixtures with cholesterol, which increases *δ* (68). *Q* is also affected by *δ* but to a lesser extent.

## Fusion of GMO Containing Formulations with Endosomal Membranes

To verify that the *Q* metric correlates to the capability of LNPs to fuse with endosomal membranes, we conducted standard FRET fusion assays. LNPs of different GMO content were prepared, and early endosomal membrane (EEM) mimetic vesicles were made comprising 40 mol % POPC, 20 mol % DOPE, 6 mol % SM, and 34 mol % cholesterol (*69*). Endosomes were also isolated from HeLa cell lines for a more biologically relevant endosome system. The sizes of these particles were measured by nanoparticle tracking analysis (NTA), shown in Table S1. The EEM vesicles and the isolated endosomes were co–labeled with 1,1’-dioctadecyl-3,3,3’,3’-tetramethylindocarbocyanine perchlorate (DiI) and 3,3’-dioctadecyloxacarbocyanine perchlorate (DiO), with DiO acting as a donor fluorophore and DiI acting as an acceptor fluorophore. The proximity of the dye molecules enables FRET to occur leading to enhanced acceptor (DiI) fluorescence. When mixed with LNPs, if EEM vesicles or isolated endosomes fuse with the LNPs, the dye molecules diffuse, increasing the distance between each other and FRET occurrences diminish. Reduced FRET allows donor (DiO) fluorescence to recover and acceptor (DiI) fluorescence to decrease. Thus, the extent of membrane fusion can be evaluated by the increase in DiO fluorescence or the decrease in DiI fluorescence (Fig. 4a).

**Figure 4:**
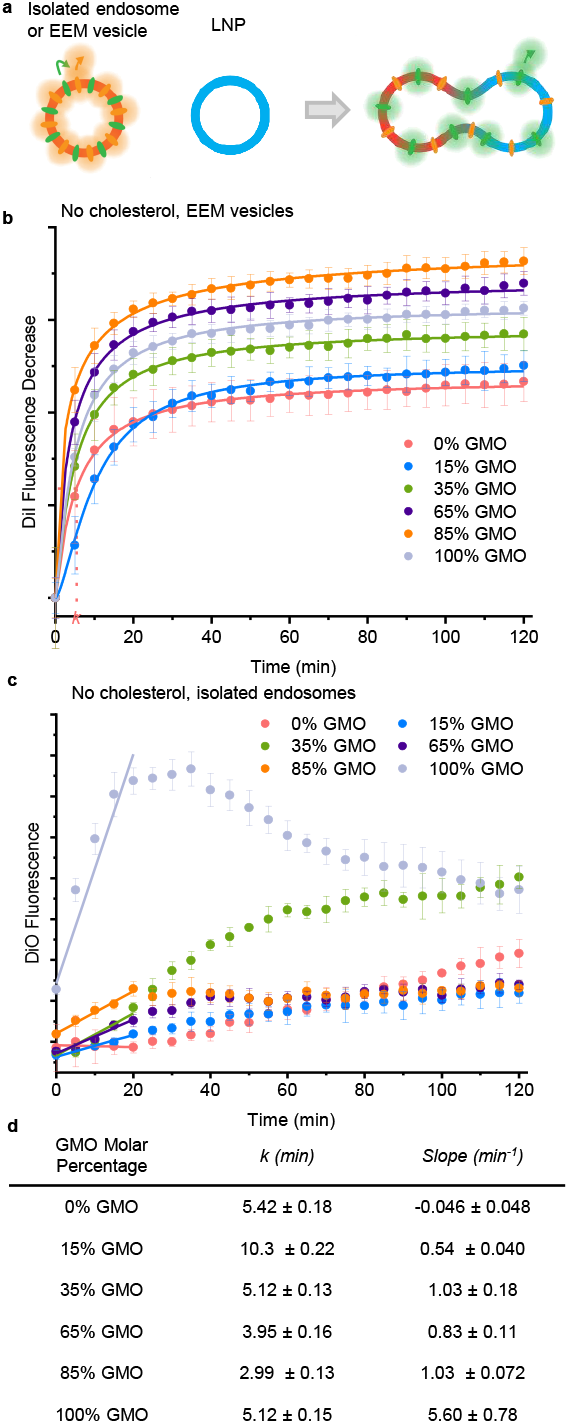
Fusion of LNPs with endosome mimics or isolated endosomes evaluated by FRET fusion assays. (**A**) Schematic of FRET assay to evaluate membrane fusion between LNPs and EEM vesicles or isolated endosomes. When membrane fusion occurs, the donor fluorescence (DiO) would increase and the acceptor fluorescence (DiI) would decrease. (**B**) Fusion assay of GMO/DOPC LNPs mixed with EEM vesicles. The DiI fluorescence decrease was measured as an indicator of fusion extent. The decrease of DiI fluorescence was fitted using a sigmoidal function where *k*, the time reaching half max fusion, can be extracted. The *k* values of different compositions are listed in (**D**). Fusion assay of GMO/DOPC LNPs mixed with isolated endosomes. The DiO fluorescence increase was measured as an indicator of fusion extent. The increase of DiO fluorescence from 0 min to 20 min was fitted linearly to extract the slope, an indicator of fusion rate. Values of the slopes of different compositions are listed in (**D**).(*n* = 3, data presented as mean ± s.d.)

For fusion with EEM vesicles, the decrease in DiI fluorescence was measured as an indicator of fusion (Fig. 4b). Over the course of 2 hours, the total fusion extent generally increased with the increase in GMO molar fraction, with the exception of 100% GMO. The DiI fluorescence decrease profiles were also fitted using a sigmoidal function to extract *k* (min), which is the time reaching half max fusion. Comparing *k* values, we can see that 85% GMO LNPs fuse with EEM vesicles the fastest, then 65% GMO and then 100% GMO or 35% GMO. According to the *Q* values we extracted, 65% GMO should be more fusogenic than 85% GMO. The reason behind the lower fusogenicity of 100% GMO LNPs in this assay is likely due to the stability of LNPs, as 100% GMO LNPs may be highly fusogenic that it fuses with each other extensively, impeding its fusion with the EEM vesicles.

The fusion assay was also performed using isolated endosomes, and in this case the increase in DiO fluorescence was measured instead (Fig. 4c). As shown in this data, the total fusion extent is highest for 100% GMO and 35% GMO and similar to the other compositions. The fusion profiles for these compositions do not fit a sigmoidal function, so to understand the fusion rate the fluorescence from 0 min to 20 min was linearly fitted instead. The slope, as listed in Fig. 4c, can be used to describe the fusion rate which is more relevant to fusogenicity since the total fusion extent can be impacted by factors such as dye diffusion rate. As seen in the slopes fitted, the fusion rate for 100% GMO is higher than 85% GMO than 65% GMO, which matches the trend we see in the *Q* values acquired.

## Structure of LNP-Endosome Fusion Intermediates

To better understand the structural attributes and morphology of the fusion processes, membrane fusion intermediates were visualized using cryo-EM. The GMO/DOPC LNPs were mixed with isolated endosomes and vitrified after 5 minutes of incubation. As shown in Fig. 5a, endosomes are vesicles with a single lipid bilayer, and many proteins are present on the endosome surface, as seen by high density of nanoscale features, or the fuzziness of the membrane. In contrast, membranes of LNPs without protein or nucleic acid moieties appear smooth and less dense. LNPs of 0% GMO (Fig. 5 b) and 35% GMO (Fig. 5 c) appear as small unilamellar vesicles and when mixed with endosomes, fusion intermediates are difficult to find. The endosomes and the LNPs are close to each other but remain separate objects. For 65% GMO (Fig. 5e) and 85% GMO (Fig. 5f) fusion intermediates are present, seen as large complex structures. To verify that the structures seen are indeed fusion intermediates, the intensity of membrane regions (highlighted by yellow lines) was line-profiled. The membrane of isolated endosomes has a characteristic peak in the middle of the membrane, as shown in the lavender region of the intensity profile, which is consistent with the fact that endosomal membranes contain transmembrane proteins such as V-ATPase proton pumps. As expected, LNP membranes do not have this characteristic peak in the membrane core region, as shown in Fig. S6,S7. Evaluating the LNP-endosome fusion structures captured in Fig. 5 e and f, it is clear that some of the membrane domains (highlighted in yellow) have a peak in the membrane core which implies that LNPs are fused with endosomal membranes. The line intensity has been profiled for different positions on the LNP-endosome fusion intermediate ultrastructure for 65% GMO (Fig. 5g) and 85% GMO (Fig. 5h), and we can see that some parts of the fused membrane regions have more endosomal content than others. LNPs with 100% GMO (no phospholipid) did not yield enough contrast due to low electron density of GMO compared to water, which makes cryo-EM imaging challenging.

**Figure 5:**
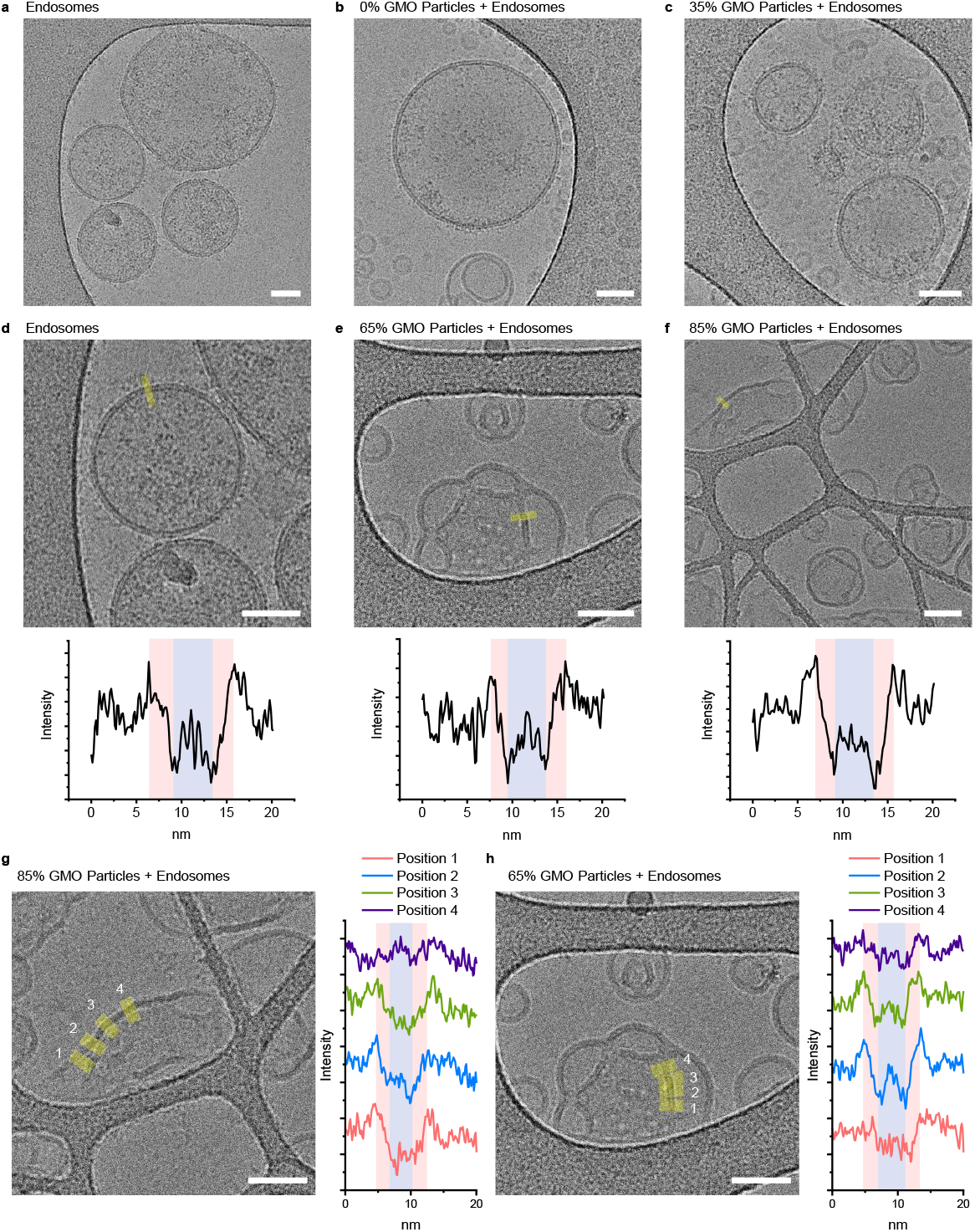
Cryo-EM micrographs of isolated endosomes mixed with GMO/DOPC LNPs and their fusion intermediates. Cryo-EM images were taken of isolated endosomes only (**A**), endosomes mixed with 0% GMO particles (**B**) and endosomes mixed with 35% GMO particles (**C**). To identify fusion intermediates, the line intensity profiles of endosome membranes were analyzed. As shown in (**D**), the intensity profile of the highlighted region in the micrograph is shown and there is a characteristic peak (shaded lavender) in the middle of the membrane region (shaded pink). Cryo-EM of a fusion intermediates from 65% GMO particles and endosomes (**E**), and 85% GMO and endosomes mixtures (**F**). The fusion intermediates are identified by a line intensity profile analysis, as the membranes show a characteristic intensity peak in the middle of membrane similar to that of endosomes.

## Fusogenicity of Ionizable Lipids

Our new platform for quantifying the fusogenicity parameter *Q* relies on measuring the change of the lattice parameters *a* of bicontinuous cubic phases as a function of temperature. For lipids such as GMO that spontaneously form bicontinuous cubic phases this is straightforward but as new LNP compounds such as ionizable lipids are synthesized, the lipid system may not necessarily have this ability. Here we show that our SAXS platform is applicable to quantify the fusogenicity of any LNP compound of interest including ionizable lipids. The method relies on measuring *a* as a function of temperature of a bicontinuous cubic phase lipid matrix of 1,2-dioleoyl-sn-glycero-3-phosphoethanolamine-N-methyl (DOPE-Me) doped with small amounts (1-5 mol %) of the lipid of interest.

As we now know, GMO increases the fusogenicity of lipid formulations so the *Q* of DOPE-Me/GMO mixtures should increase as GMO content increases. DOPE-Me/GMO mixtures were made and SAXS measurements were done at temperatures 50-90°C at 5 °C increments. The *M* values were then extracted (Table 4 and Fig. 6a,b), and we can see that with increasing amounts of GMO *M* becomes more negative. From Table 4 and Fig. 6, it is evident that *Q* increases with increasing amounts of GMO, implying that GMO makes DOPE-Me membranes more fusogenic, which is consistent with our previous findings that elevating GMO content increases fusogenicity. Therfore, DOPE-Me can be used as host matrix for evaluating the fusogenicity of exogenous lipids. We then used this method to study the fusogenicity of ionizable lipids commonly used in LNP systems, such as SM-102 and ALC-0315 which were respectively used in the Moderna and Pfizer/BioNTech COVID-19 vaccines. Samples were prepared in both water as well as sodium acetate (NaAc) buffer at pH 4, which represents the acidic switch of the endosomal environment. Small amounts of ionizable lipids (1.5% and 5%) were included in the DOPE-Me host membrane, and the lattice parameter change with temperature was measured using SAXS. Figure 6 d shows that the *M* of DOPE-Me shifts with the inclusion of ionizable lipids, and interestingly *M* becomes more positive, which is the opposite trend of GMO. We find that *Q* becomes more positive with the inclusion of IL, which indicates that these ionizable lipids increase the fusogenicity of DOPE-Me (Table 5). The values extracted were from the *Im*3*m* phase as it is the dominant cubic phase in these compositions. It is noteworthy that SM-102 is more fusogenic than ALC-0315 as *Q* is significantly higher when compared to the DOPE-Me matrix. In contrast, DSPC (1,2-Distearoyl-sn-glycero-3-phosphocholine) which is commonly used in LNP formulations as a stabilizer lipid, has a very small effect on *Q* of DOPE-Me. This result supports that ionizable lipid compounds of LNP formulations are the major driver of membrane fusion and endosomal escape. The difference in *Q* of DOPE-Me with various added lipids further indicates the robustness of this fusogenicity quantification platform, as different lipids shift the *Q* of DOPE-Me to a distinct extent depending on their fusion capability.

**Table 4:** The 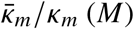 values extracted and *Q* calculated for DOPE-Me with increasing amounts of GMO at 37°C.

| DOPE-Me/GMO Molar Ratio | Phase | $M$ | $Q$ |
| --- | --- | --- | --- |
| 98.5/1.5 | *Im3m* | $-0.849 \pm 0.034$ | $0.004 \pm 0.032$ |
| 98.5/1.5 | *Pn3m* | $-0.854 \pm 0.031$ | $-0.001 \pm 0.028$ |
| 97/3 | *Im3m* | $-0.845 \pm 0.018$ | $0.019 \pm 0.016$ |
| 97/3 | *Pn3m* | $-0.848 \pm 0.011$ | $0.015 \pm 0.009$ |
| 95/5 | *Im3m* | $-0.851 \pm 0.012$ | $0.027 \pm 0.010$ |
| 95/5 | *Pn3m* | $-0.874 \pm 0.008$ | $0.004 \pm 0.005$ |
| 90/10 | *Pn3m* | $-0.881 \pm 0.005$ | $0.034 \pm 0.002$ |

**Table 5:** The 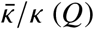 values extracted for DOPE-Me with small amounts of commonly used lipid nanoparticle lipids with different buffers at 37°C.

| Buffer |  | H<sub>2</sub>O |  |
| --- | --- | --- | --- |
| Added Lipid | DSPC | ALC-0315 | SM-102 |
| 0% |  | 0.006 ± 0.003 |  |
| 1.5% | 0.030 ± 0.004 | -0.045 ± 0.002 | 0.038 ± 0.001 |
| 5% | -0.032 ± 0.006 | 0.131 ± 0.003 | 0.179 ± 0.012 |

| Buffer |  | NaAc buffer |  |
| --- | --- | --- | --- |
| Added Lipid | DSPC | ALC-0315 | SM-102 |
| 0% |  | 0.031 ± 0.013 |  |
| 1.5% | -0.023 ± 0.004 | -0.025 ± 0.006 | -0.045 ± 0.009 |
| 5% | 0.012 ± 0.018 | 0.131 ± 0.001 | 0.151 ± 0.001 |

**Figure 6:**
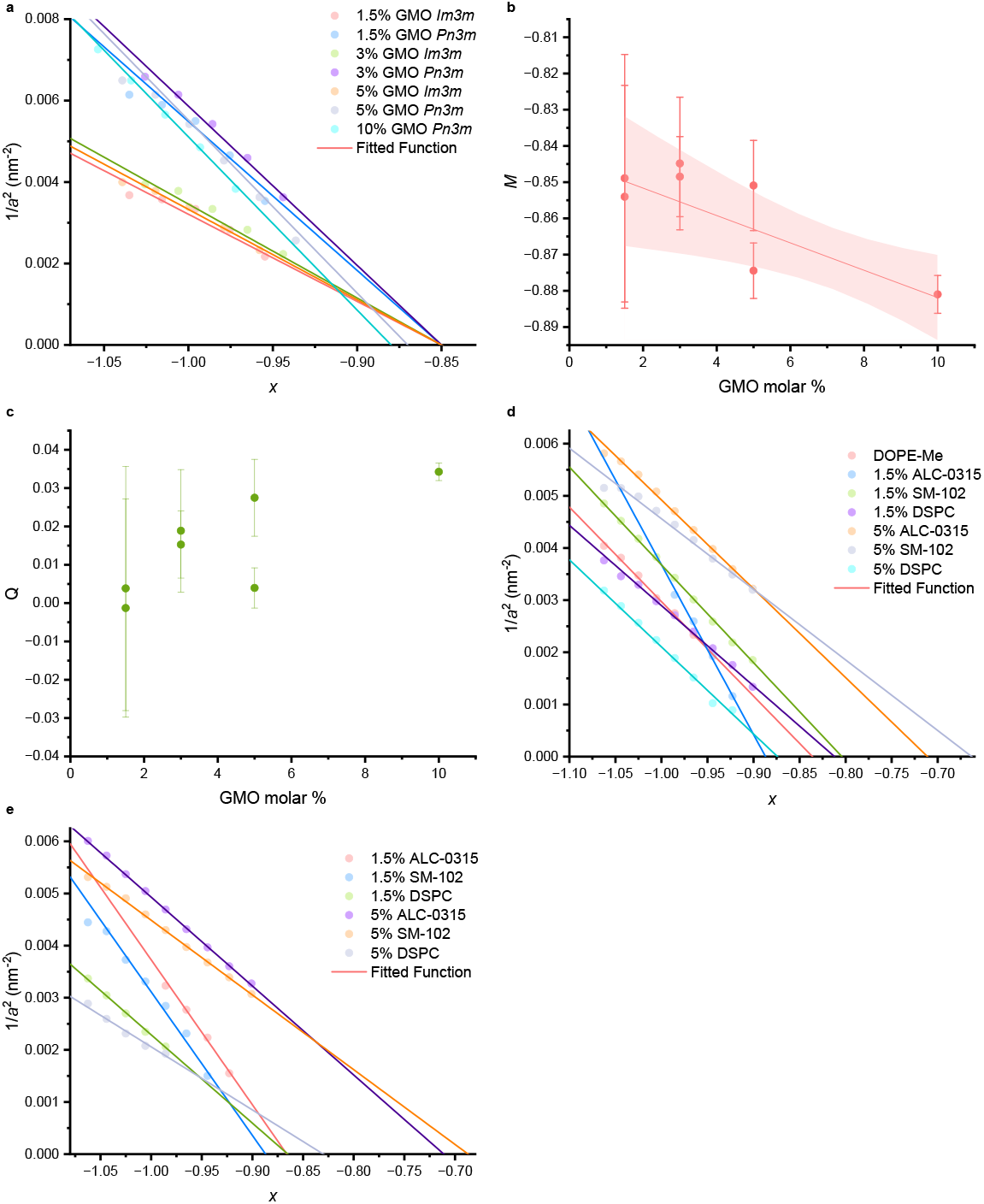
Fusogenicity of lipids are evaluated by the change in *Q* of DOPE-Me. (**A**)Plot of 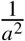 against *x* for DOPE-Me with increasing amounts of GMO, with the fitted linear function where the x intercept is *M. M* (**B**) and *Q* (**C**) plotted against GMO molar percentages. *M* becomes more negative as GMO content increases. The red line is a linear regression fit with the 95% confidence interval shown (red shade). *Q* increases as GMO content increases. (**D**,**E**) Plots of 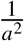 against *x* for DOPE-Me with 1.5 % and 5 % of ionizable lipid ALC-0315, SM-102 and helper lipid DSPC in water (**D**) or NaAc buffer (**E**).

## Molecular dynamics simulations

To corroborate our experimental results which suggest that *Q* is an important determinant of membrane fusion, we turn to coarse-grained molecular dynamics. A distinct advantage here is that we can continuously tune the elastic parameters, most importantly *Q*, via temperature and lipid shape.

In a recent work, we have proposed a new computational method to measure 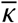 from the energy of TPMS unit cells in a highly coarse-grained lipid model. There, we measured 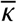 over a range of temperatures and lipid shapes. Here, over the same range, we simulate multiple vesicle-planar fusion events for each system. Figure 7 shows the probability of fusion and long-lived hemifusion, pooled together, as a function of different elastic parameters. The probability shows the strongest dependence on the bilayer elastic ratio 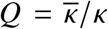, even more so than the value of 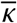 relative to the thermal energy scale.

**Figure 7:**
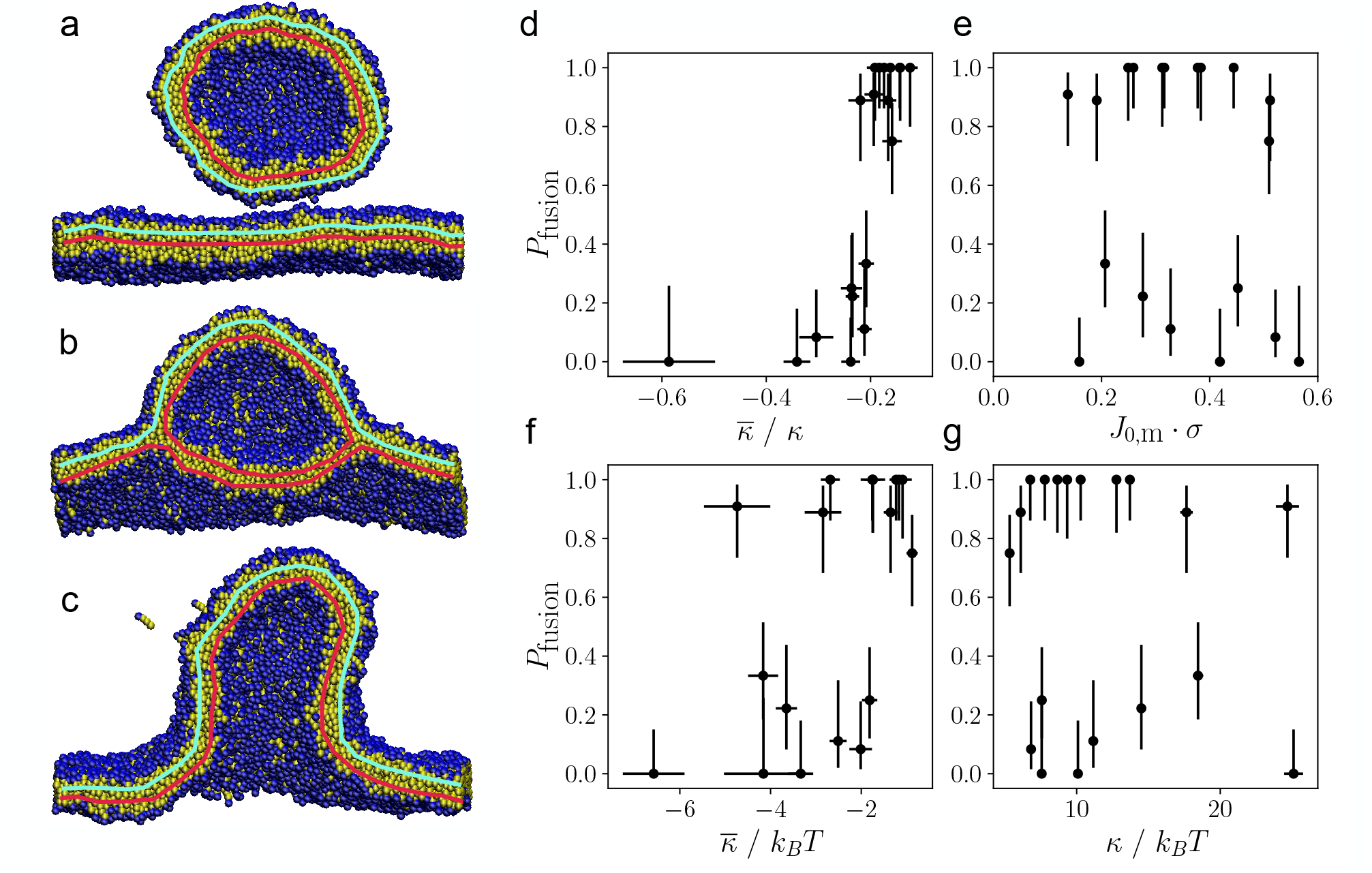
Fusogenicity of lipids as correlated with various elastic parameters in coarse-grained molecular dynamics simulations. (**a-c**) Snapshots of simulations showing unfused, hemifused, and fused states of a vesicle approaching a plane. The cis- and trans-leaflets in each case are indicated with cyan and red, respectively. (**d-g**) Plots of the probability that fusion or hemifusion will occur versus the bilayer elastic ratio 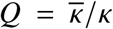, the spontaneous monolayer curvature *J*_0,m_, the absolute bilayer Gaussian modulus 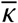, and the ordinary bending rigidity *κ*. Each data point is calculated from the result of 6-12 simulations. Error bars indicate the Clopper-Pearson interval at 67% confidence.

Intriguingly, the spontaneous monolayer curvature *J*_0_ as calculated by stress profile measurements seems entirely uncorrelated with the probability of fusion. Actually, a convenient quirk of the implicit solvent model we use is that the dependence of *J*_0_ on *T* subverts expectations, *i*.*e*. here *J*_0_ increases with increased *T* . Since 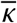 nevertheless still softens with *T*, this allows us to disentangle the separate effects of 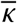 and *J*_0_ on fusion.

### Discussion

Despite the success in LNP mediated drug delivery as exemplified with the COVID mRNA vaccine, endosomal escape is still a major challenge in achieving efficient LNP drug delivery. Most LNPs developed currently use ionizable lipids to induce endosomal escape, which depends on the electrostatic interaction between the positively charged ionizable lipid and negatively charged endosomal membrane, as well as inducing phase transition to an inverse hexagonal phase. However, designing more fusogenic LNPs has been difficult due to lacking a quantitative measure for lipid formulation fusogenicity. In this study, we propose a parameter *Q* that quantifies the fusogenicity of LNPs by extracting the 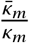 value from lipid compositions that form bicontinuous cubic phases. The 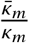 and *Q* were successfully determined for GMO/DOPC and GMO/DOPC/Chol lipid formulations, and their fusion with EEM vesicles and isolated endosomes were evaluated and shown to match the trend in *Q*.

To evaluate the fusogenicity of lipids commonly used in LNPs that do not necessarily form cubic phases, we used DOPE-Me as a host lipid matrix for cubic phases and added small amounts of ionizable lipids or helper lipids. We observed that *Q* of DOPE-Me changes with the inclusion of ILs, and that ILs used in FDA-approved formulations such as SM-102 and ALC-0315 increased *Q* significantly. In both water and acidic buffer, SM-102/DOPE-Me mixtures have a higher *Q* compared to that of ALC-0315/DOPE-Me mixtures, which shows that SM-102 is more fusogenic than ALC-0315. On the contrary, DSPC, a commonly used helper lipid in LNP formulations, has a much lower impact on *Q* of DOPE-Me. As a result DSPC have low fusogenicity compared to the IL compounds. Our molecular dynamic results have also shown that the probability of fusion and hemifusion shows the strongest dependence on *Q* among the different elastic parameters.

These results show how our platform establishing the new parameter *Q* can successfully evaluate the fusogenicity of different lipids. In the future, *Q* can be measured to guide ionizable lipid design. Effective ionizable lipids can be tested for fusogenicity and when fusogencitiy is low, lipids can be optimized to be more fusogenic, and when fusogenicity is high other factors of LNPs can be targeted such as size and stability. Previously researchers have lacked information on which physical parameters of lipids influence LNP biological performance, and our platform provides an important physical parameter that can guide in the engineering of more effective LNPs. Moreover, this platform can be used to work towards a more comprehensive understanding of membrane fusion, which is an important component of biological systems. It is also worth noting that this platform may also work for quantifying the fusogenicity of peptides in lipid membranes, which is a direction that warrants further investigation and could expand our platform to not only lipids but a variety of biologically relevant molecules.

## Acknowledgments

This research was in part conducted in the Materials Research Laboratory in University of Illinois Urbana-Champaign. We thank the beamline scientists at Advanced Photon Source, beamline 12-ID-B, Argonne National Laboratory. We thank Dr. Alexander J. Sodt at National Institutes of Health for discussion regarding the sign of monolayer Gaussian modulus.

## Author contributions

L.Z. and C.L. designed research; L.Z., M.B., S.G., A.G., J.R., and Y.G performed research; L.Z., S.G., M.D. and C.L. analyzed data; and L.Z., S.G. and C.L. wrote the paper.

## Data and materials availability

All data needed to evaluate the conclusions in the paper are present in the paper and/or the Supplementary Materials.

## Supplementary materials

Materials and Methods

Supplementary Text

Figs. S1 to S12

Tables S1 to S2

References *(7-71)*

