## Supplementary Materials for "Introducing a fusogenicity metric for lipid nanoparticle formulation"

**This PDF file includes:**

Materials and Methods

Supplementary Text

Figures S1 to S12

Tables S1 to S2

### Materials and Methods

#### Materials

Glycerol monoooleate (GMO) and cholesterol (Chol) were purchased from Sigma-Aldrich (MO, USA). 1,2-dioleoyl-*sn*-glycero-3- phosphocholine (DOPC), 1,2-dioleoyl-*sn*-glycero-3-phosphoethanolamine-N-[methoxy (polyethylene glycol) - 2000] (DOPE-PEG), 1,2-dioleyl-sn-glycero-3-phosphoethamolamine (DOPE), 1-hexadecanoyl-2-(9Z-octadecenoyl)-sn-glycero-3-phosphocholine (POPC) and sphingomyelin from porcine brain (SM) were purchased from Avanti Polar Lipids (AL, USA). 1,1'-dioctadecyl-3,3,3',3'- tetramethylindocarbocyanine perchlorate (DiI) and 3,3'-dioctadecyloxacarbocyanine perchlorate (DiO) were purchased from Biotium (CA, USA).

#### Small angle X-ray scattering (SAXS)

Lipid formulations (total lipid concentration 200 mM) were prepared. Lipid chloroform solutions were mixed at desired volumetric ratios. Chloroform was removed by treating the solution with a stream of nitrogen and then placing it under vacuum for at least 8 hrs. The dried lipid film was hydrated and incubated overnight and transferred to quartz capillaries (Hilgenberg Glas, Germany). Synchrotron SAXS were performed at beamline 12-ID-B of the Advanced Photon Source at Argonne National Laboratory. The average photon energy was 14 keV and the data was radially averaged upon acquisition on a Pilatus 2M detector.

For DOPE-Me with addition of GMO experiments, SAXS measurements were collected with in-house equipment composed of a Xenocs GeniX3D Cu K$\alpha$ ultralow divergence X-ray source (1.54Å\ 8 keV), with a divergence of ~1.3 mrad. The 2D diffraction data were radially averaged upon acquisition on a Pilatus 300 K 20 Hz hybrid pixel detector (Dectris).

BioXTAS RAW (*70*) and Origin 2025 (OriginLab, Inc.) were used to process the SAXS data, and in-house Python scripts were used to extract the $M$ and $b$ values using equation 2.

#### Preparation of LNPs

LNPs were formed by NanoAssemblr Ignite (Precision NanoSystems). Lipid chloroform solutions were mixed at desired volumetric ratios and the solvent is removed as described above. The lipids were then dissolved in ethanol at a concentration of 10 mM. Total flow rate was maintained at 12 mL per min for all formulations. A flow rate ratio of 4:1 (aqueous to ethanol) was used. Ethanol was removed by dialysis using Slide-A-Lyzer™ (ThermoFisher) cassettes with a molecular weight cutoff of 3.5k. Particle concentration and size were measured by nanoparticle tracking analysis with Nanosight NS300 (Malvern Panalytical).

**Membrane fusion studies**

The EEM comprised 40 mol % POPC, 20 mol % DOPE, 6 mol % SM, and 34 mol % cholesterol (*69*). 0.1 mol % DiO and 0.1 mol % DiI were included for FRET labeling. Endosomes were isolated with Trident endosome isolation kit (GeneTex). Particle concentration and size were measured by nanoparticle tracking analysis with Nanosight NS300 (Malvern Panalytical).

For FRET assay, endosomes were incubated with 20 $\mu$M DiO and 20 $\mu$M DiI for 1 hr at 37 °C for colabeling. Free dye was removed and endosomes were resuspended in PBS. Endosomes and lipid-siRNA complexes were incubated at 37°C and measured for DiO fluorescence every 5 minutes. Fluorescence was detected with Synergy Neo 2 microplate reader (Biotek).

**Cryogenic electron microscopy**

To prepare samples for cryo-EM imaging, holey carbon coated 300 mesh copper grids (Electron Microscopy Sciences) were glow discharged at 15 mA for 30 s with PELCO easiGLOW™ glow discharge system (Ted Pella). 2 $\mu$L of LNPs and 2 $\mu$L of endosomes was applied to the grids and incubated for 10 min. The grids were then blotted for 2.5 s and plunge frozen in liquid ethane using Vitrobot Mark IV, under 4 °C and 100% humidity. The grids were kept in liquid nitrogen until imaging. Cryo-EM images were collected with Glacios Cryo-TEM (ThermoFisher) at 200 kV with a Falcon 4 direct electron detector. Images were taken at -3 $\mu$m defocus to improve contrast.

**Molecular dynamics simulations**

**Setup**

The coarse-grained model used and how lipid shape is modified is described in our recent work currently in review. Approximately 4,000 and 8,000 lipids make up the spherical vesicle and the flat bilayer, respectively. The flat bilayer, which is periodically connected over its plane, is kept under zero tension. The box size in the up-down direction is kept fixed. A small downward force is applied to each bead of the vesicle and a proportional upward force is applied to each bead of the plane such that they are gently brought into contact.

For the simulation engine, we use the espresso package. A line in the espresso source code must be changed to keep thermal fluctuations in the fixed up-down direction.

**Analysis**

The new method of measuring $\overline{\kappa}$ is described in detail in our recent work currently in review. We use the heuristic joint fits there to infer values of $\overline{\kappa}$ and $\overline{\kappa}/\kappa$ for those systems where an explicit calculation was not made. The method of measuring the bending rigidity by shape fluctuations is outlined in the same paper.

The spontaneous monolayer curvature $J_0$ is obtained by the first moment of the lateral stress profile $\sigma(z)$ of a tension-less leaflet:

$$-\kappa_{\mathrm{m}} J_0 = \int_0^{\infty} \mathrm{d}z \sigma(z) z \tag{S1}$$

Measured about the midplane of a flat bilayer with 256 lipids. Again, as with the other elastic parameters, we use a heuristic joint fit to infer the values of J0 where an explicit measurement was not made.

For the fusion simulations, fusion (or hemifusion) is identified when the number of leaflets drops below four. The number of leaflets is identified by a single linkage clustering algorithm where two lipids are identified with the same leaflet if their distance is less than 3 CG units and the cosine similarity of their directors is greater than 0.9. We do not differentiate between fused and hemifused states, counting both as fusion here. In principle, we could separately identify the fused and hemifused states as having two and three leaflets separately. However, the wide variety of different intermediates lead to a noisy signal in the number of leaflets identified with the described method.

### Supplementary Text

#### Expression of b

$b$ from equation 2 is expressed as the following, derived by Siegel et.al. from (*34*).

$$b = -\frac{\kappa_2}{\kappa_m}\frac{S_2}{S_1} \tag{S2}$$

$S_N$ is a dimensionless coefficient of cubic phases, which differs depending on the space group. $S_N = \int\int K^N dA$, where $K$ and $dA$ are defined on the minimal surface.

#### Monolayer to bilayer

The relation between the monolayer and bilayer Gaussian curvature modulus is obtained by constructing a bilayer free energy out of two monolayer free energies and identifying the term proportional to $K$ as the bilayer modulus $\overline{\kappa}$. However, there is a discrepancy in the literature regarding the final form of this relation due to a subtle identification of the reference state. Since we pick a side, it is worth presenting why. Here we will closely follow the derivation in the Appendix of (*34*) while being slightly more explicit.

The monolayer curvature free energy density for the upper (+) and lower (-) leaflets of a symmetric bilayer are:

$$f_{\text{bend},\pm} = \frac{1}{2}\kappa_{\text{m}}(J_\pm \mp J_0)^2 + \overline{\kappa}_{\text{m}}K_\pm \tag{S3}$$

We additionally can include a leaflet stretching free energy density which penalizes deviations from the rest area $A_0$ of the leaflets by a compressibility modulus $k_{A,\text{m}}$:

$$f_{\text{st},\pm} = \frac{1}{2}k_{A,\text{m}}\frac{(A_\pm - A_0)^2}{AA_0} \tag{S4}$$

Note that the density is taken as free energy per *midplane*[] area A.

For simplicity, and without loss of generality for the purposes of calculating $\overline{\kappa}$, we will consider that the bilayer midplane is a minimal surface $J = 0$. Then, we can make the following identifications for parallel surfaces displaced from the midplane by a signed displacement $\delta$:

$$\mathrm{d}A_\pm = (1 + K\delta^2)\mathrm{d}A \tag{S5}$$

$$A_\pm = A + \delta^2 \int K\mathrm{d}A \tag{S6}$$

$$J_\pm = \frac{2K\delta}{1 + K\delta^2} \tag{S7}$$

$$K_\pm = \frac{K}{1 + K\delta^2} \tag{S8}$$

The bilayer free energy density reads:

$$f_\mathrm{B} = 2f_\mathrm{st} + \frac{\mathrm{d}A_\pm}{\mathrm{d}A}\left(f_{\mathrm{bend},+} + f_{\mathrm{bend},-}\right) \tag{S9}$$

$$= 2f_\mathrm{st} + (1 + K\delta^2)\left(f_{\mathrm{bend},+} + f_{\mathrm{bend},-}\right) \tag{S10}$$

Which can be expanded for small $K$ after the appropriate substitutions for the geometry of the monolayers as:

$$f_\mathrm{B} = k_0 + k_1 K + O(K^2) \tag{S11}$$

Where we identify

$$k_0 = 2f_\mathrm{st} + \kappa_\mathrm{m} J_0^2 \tag{S12}$$

$$k_1 = 2(\overline{\kappa}_\mathrm{m} - 2\kappa_\mathrm{m}\delta J_0 + \frac{1}{2}\kappa_\mathrm{m}(\delta J_0)^2) \tag{S13}$$

The last expression can be found in the literature with the final term $\sim \delta^2$ omitted. Where does this discrepancy come from? If we had instead written the erroneous expressions for a monolayer and bilayer free energy density as:

$$f'_\pm = \sigma_\pm + f_{\mathrm{bend},\pm} \tag{S14}$$

$$f'_\mathrm{B} = \frac{\mathrm{d}A_\pm}{\mathrm{d}A}\left(f'_+ + f'_-\right) \tag{S15}$$

Where $\sigma_\pm$ are the leaflet tensions, then we would get the expansion coefficients:

$$k'_0 = \kappa_\mathrm{m} J_0^2 + \sigma_+ + \sigma_- \tag{S16}$$

$$k'_1 = 2(\overline{\kappa}_\mathrm{m} - 2\kappa_\mathrm{m}\delta J_0 + \frac{1}{2}\delta^2(\kappa_\mathrm{m} J_0^2 + \sigma_+ + \sigma_-)) \tag{S17}$$

Requiring $k_0'$ to be zero then leads to the quadratic order term in $k_1'$ disappearing. However, the tension $\sigma_\pm$ is not the stretching free energy density. The stretching free energy density is instead given by the work required to stretch the membrane from its rest value $A_0$:

$$f_{\mathrm{st},\pm}(A) = \frac{1}{A}\int_{A_0}^{A} \mathrm{d}A' \sigma_\pm(A') \tag{S18}$$

where, if we write $\sigma_\pm(A)$ at linear order in area strain as $\sigma_\pm(A) = k_A \frac{A-A_0}{A_0}$, we recover the stretching free energy density used above wherein the individual leaflet tensions do not appear in the expression for the Gaussian modulus.

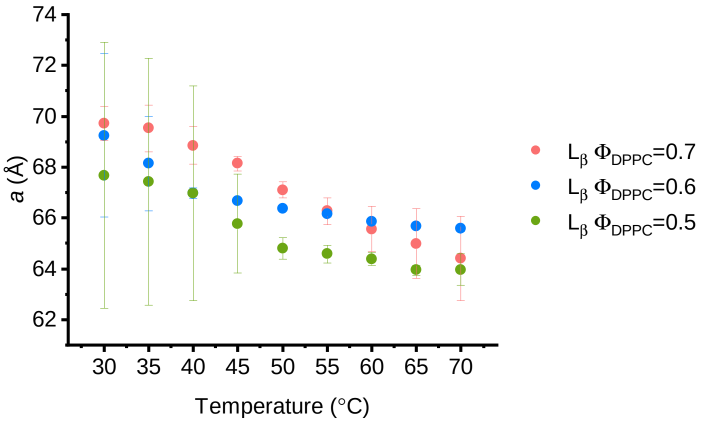


**Figure S1**: **The lattice parameters of lamellar ($L$) phases of the DPPC/Chol compositions at different temperatures determined by SAXS.** ($n$ = 3, data presented as mean ± s.d.)

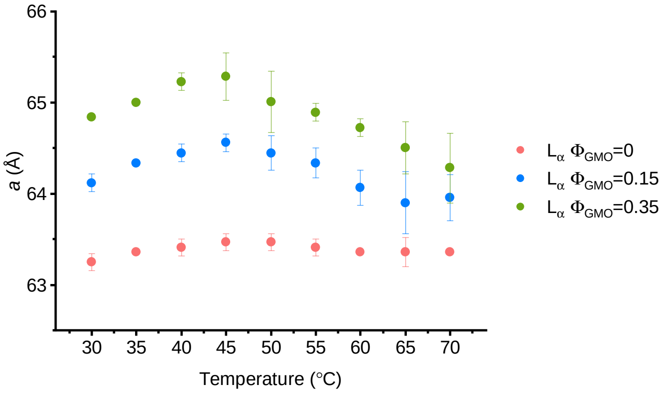


**Figure S2**: **The lattice parameters of lamellar ($L$) phases of the GMO/DOPC compositions at different temperatures determined by SAXS.** ($n$ = 3, data presented as mean ± s.d.)

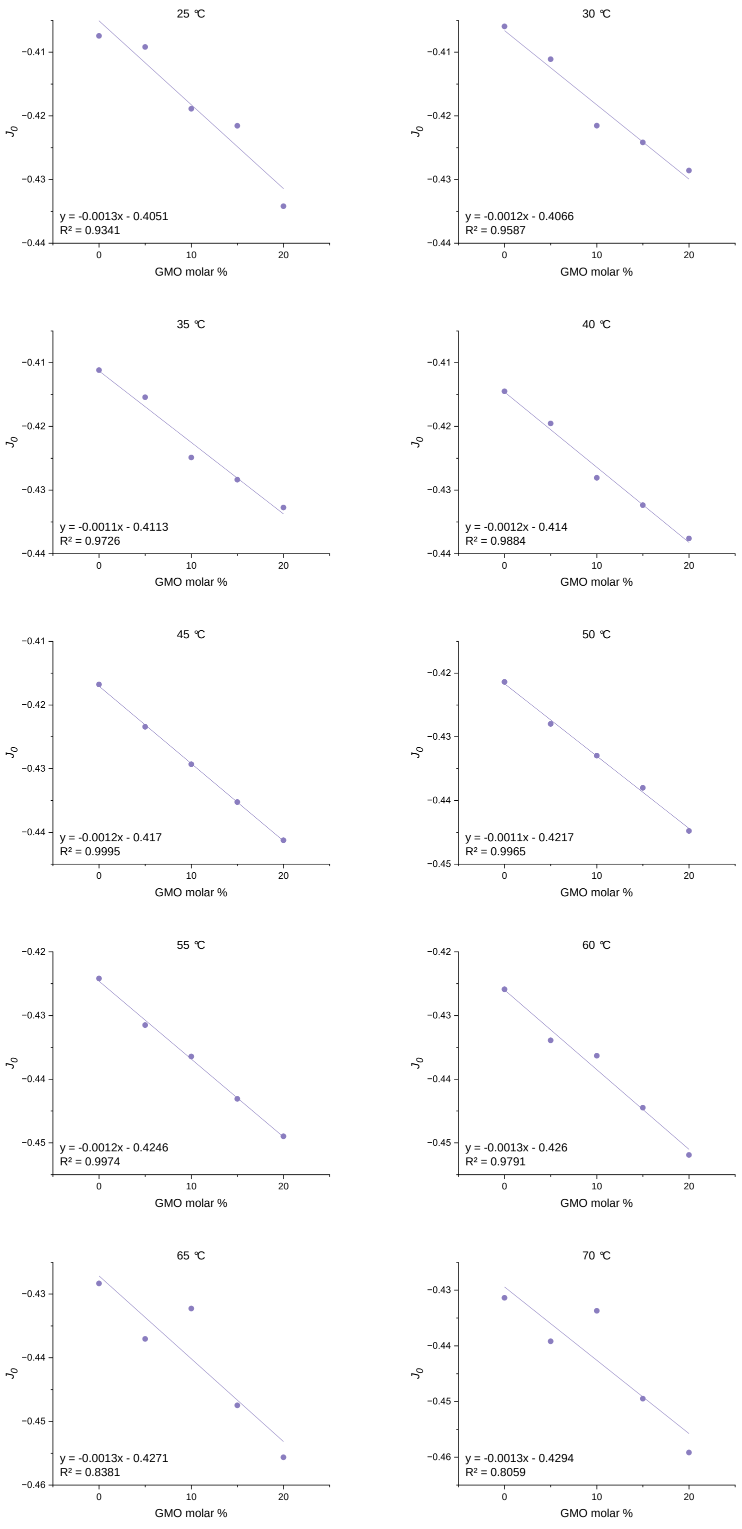


**Figure S3**: **Calculation of $J_0$ of GMO at different temperatures.** Calculated $J_0$ ($nm^{-1}$) value of each DOPE/GMO mixture plotted against GMO molar percentage at different temperatures. The $J_0$ of GMO is obtained from the extrapolation of the linear fitting at molar percentage of GMO equal to 100%.

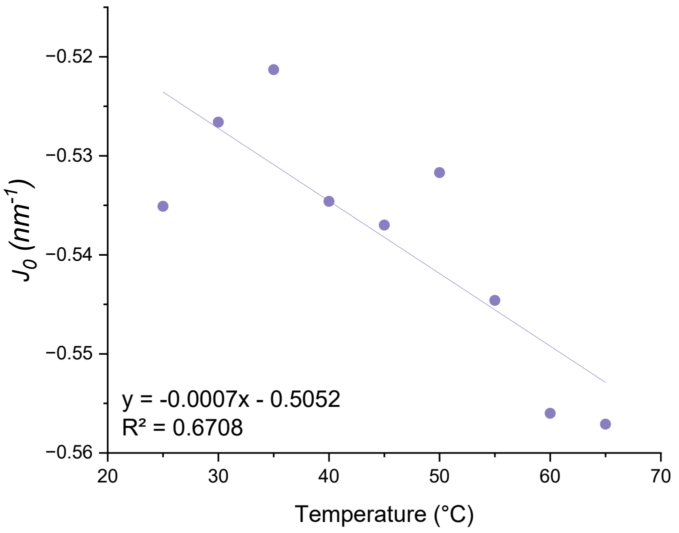


**Figure S4**: $J_0$ **of GMO dependence on temperatures.** Calculated $J_0$ (nm$^{-1}$) value of GMO obtained from Figure S3 is plotted against temperature. The relationship of GMO $J_0$ with temperature is obtained from linear fitting.

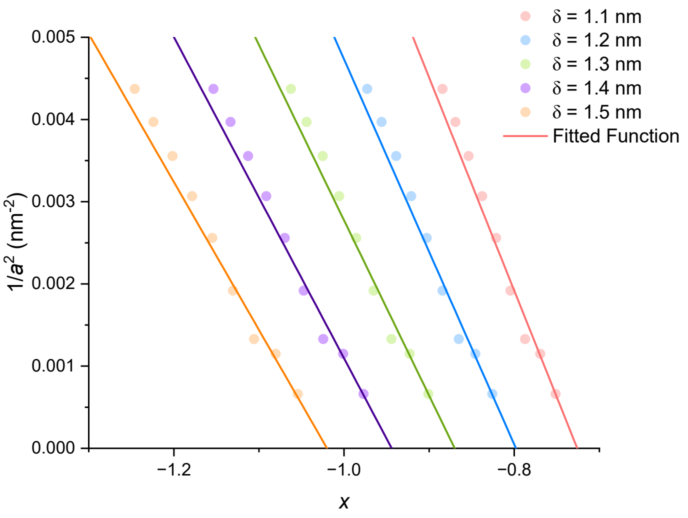


**Figure S5**: **Plots of** $\frac{1}{a^2}$ **against** $x$ **for DOPE-Me compositions using different values of** $\delta$**.**The fitted linear function where the x intercept is $M$.

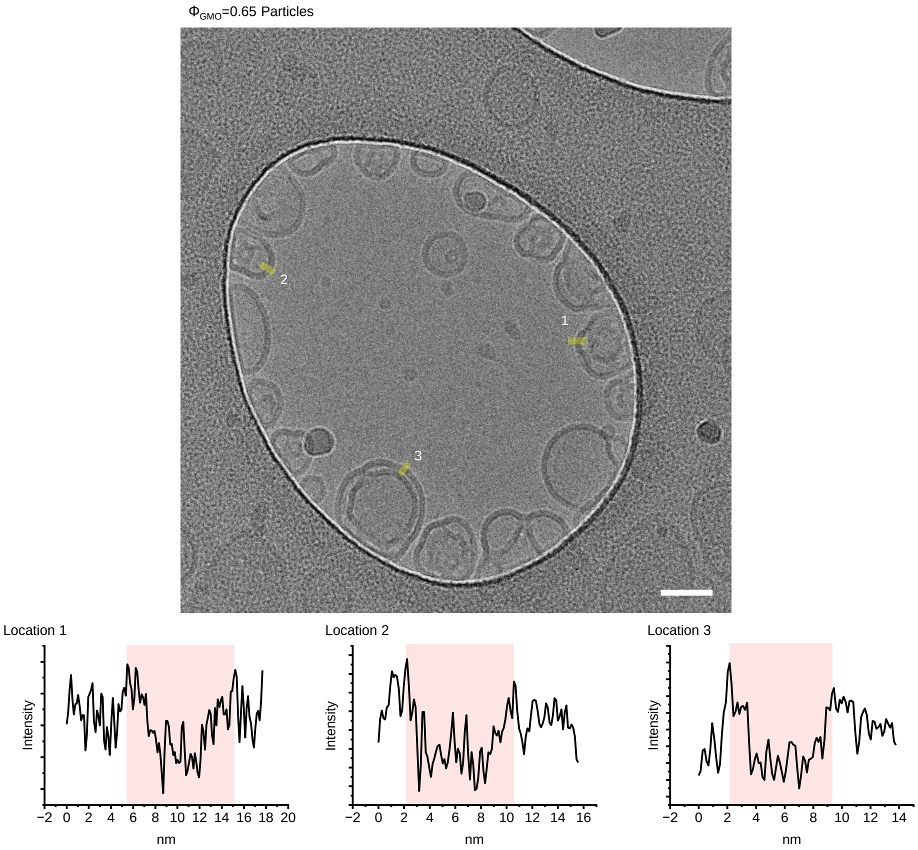


**Figure S6**: **Cryo-EM micrographs of 65% GMO particles.** Line intensity profile analysis was done on the highlighted regions, and none of them has the characteristic intensity peak in the middle of the membrane as endosomal membranes do.

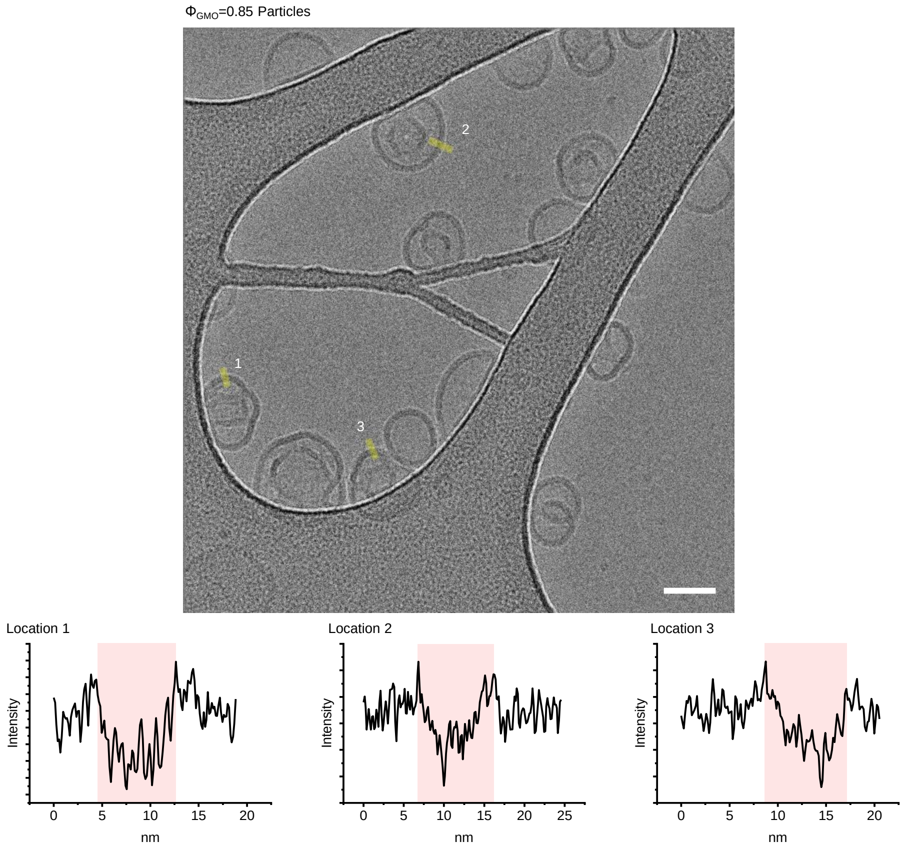


**Figure S7**: **Cryo-EM micrographs of 85% GMO particles.** Similar to Figure S6, and the characteristic endosomal membrane intensity peak is not present.

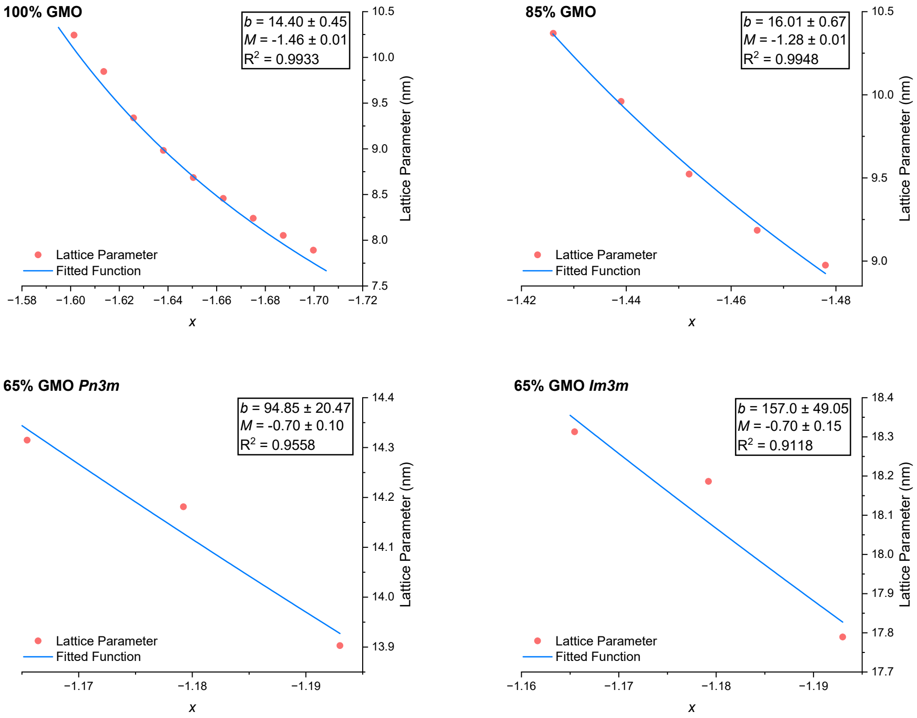


**Figure S8**: **Fitting for *M* of GMO/DOPC Mixtures.** *M* of GMO/DOPC mixtures is fitted with equation 2 using the bicontinuous cubic phase lattice parameters measured. The $x$ values are calculated from equation 3, using the spontaneous curvature of the mixtures at the corresponding temperature and a $\delta$ value of 1.3 nm.

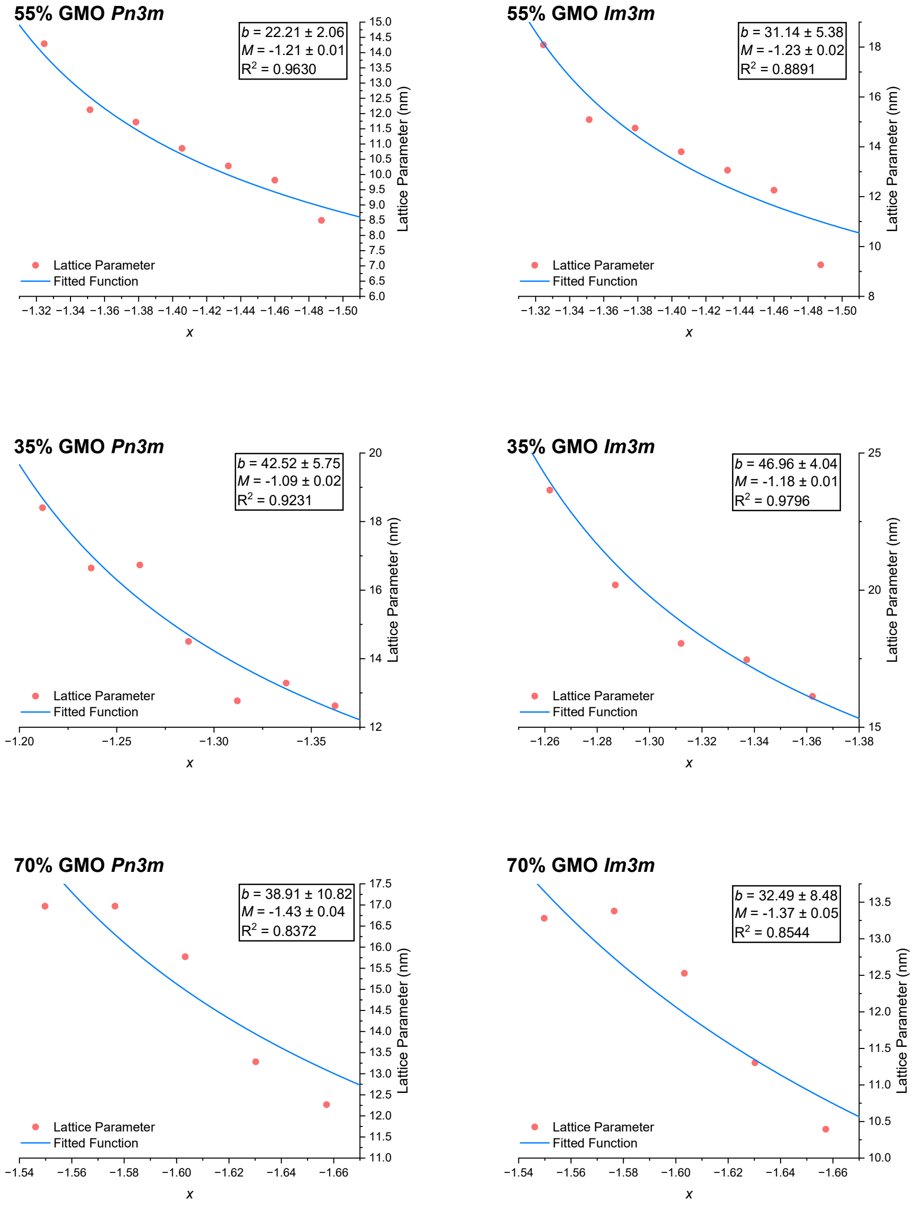


**Figure S9**: **Fitting for $M$ of GMO/DOPC/Chol Mixtures.** $M$ of GMO/DOPC/Chol mixtures is fitted in the same way as Figure S8.

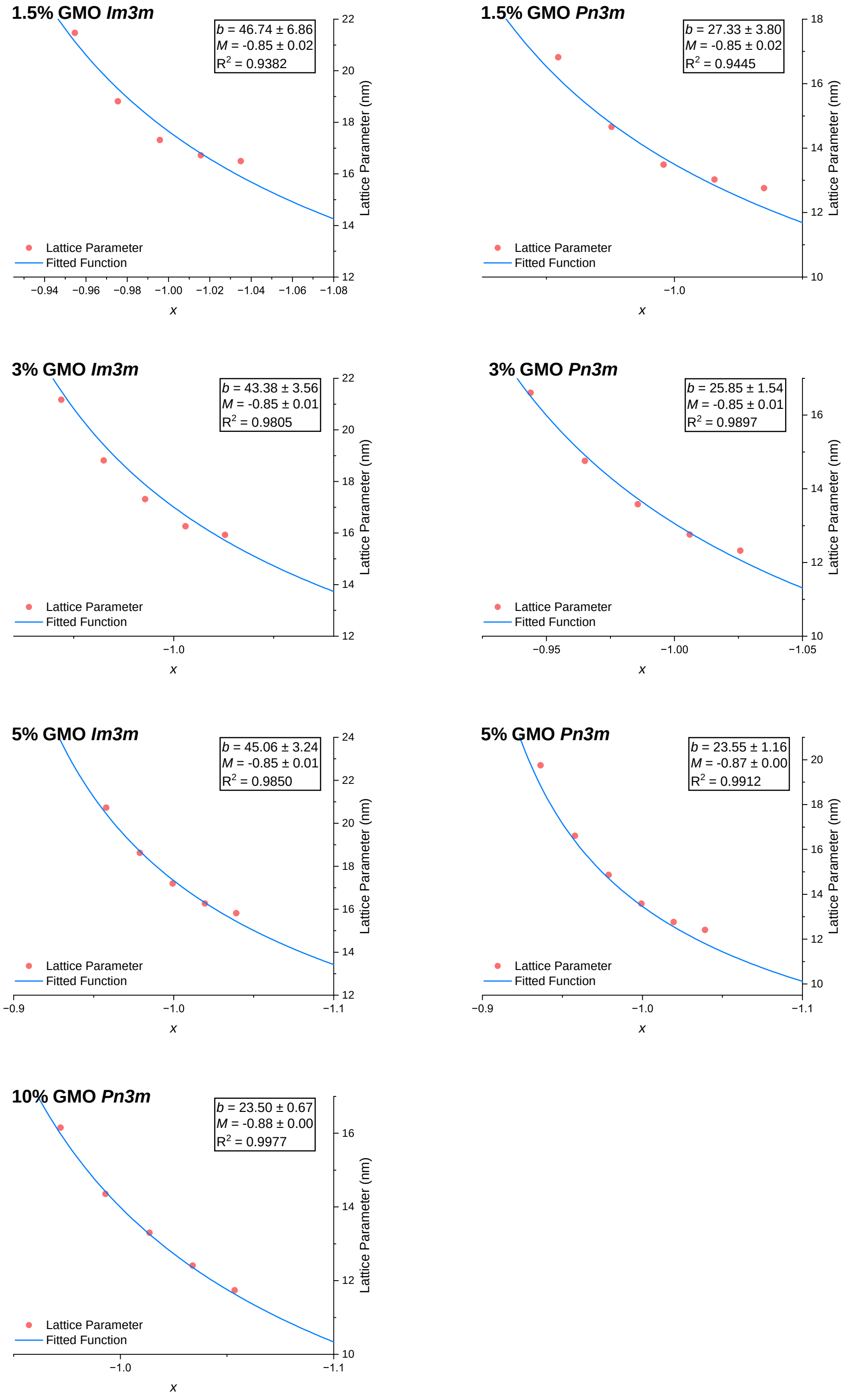


**Figure S10**: **Fitting for** $M$ **of DOPE-Me/GMO Mixtures.** $M$ of DOPE-Me GMO mixtures is fitted in the same way as Figure S8.

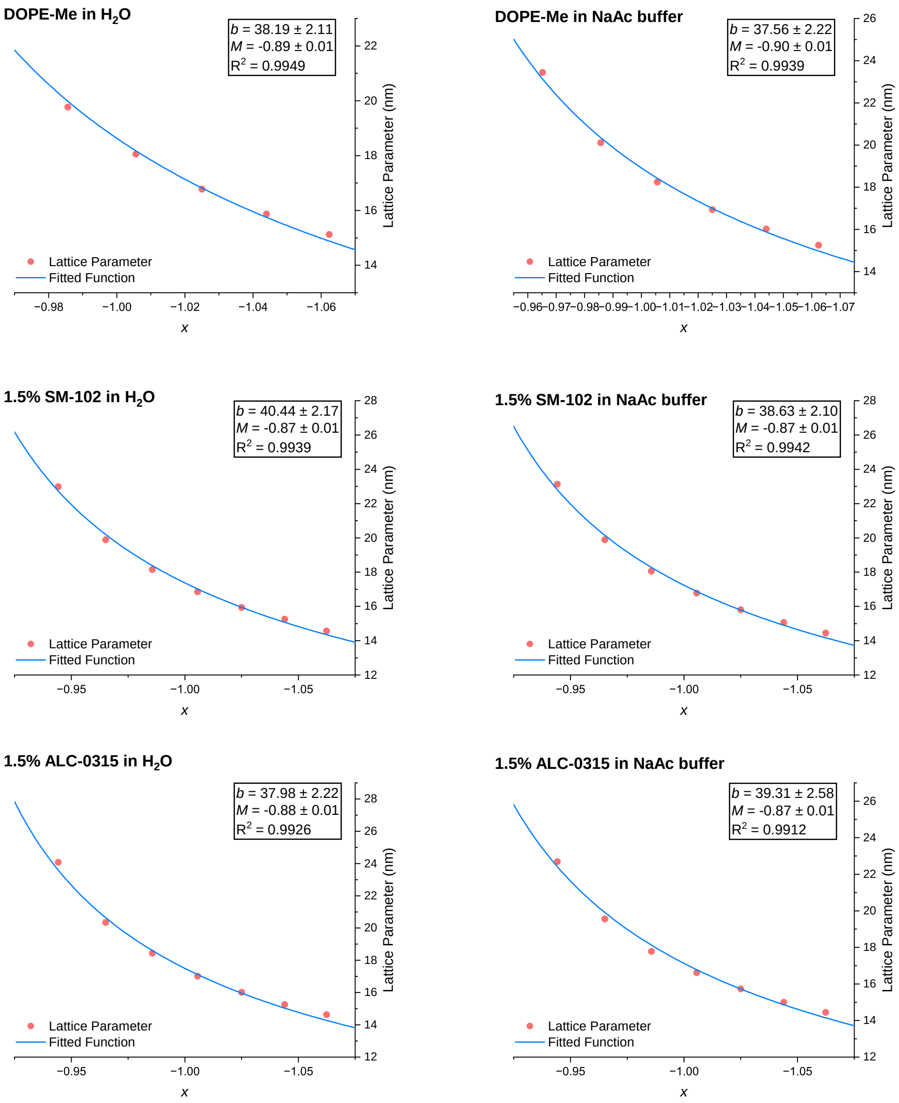


**Figure S11**: **Fitting for $M$ of DOPE-Me/IL Mixtures.** $M$ of DOPE-Me IL mixtures is fitted in the same way as Figure S8. The IL used are SM-102 and ALC-0315, which are lipids used in the COVID-19 vaccine formulation (*71*).

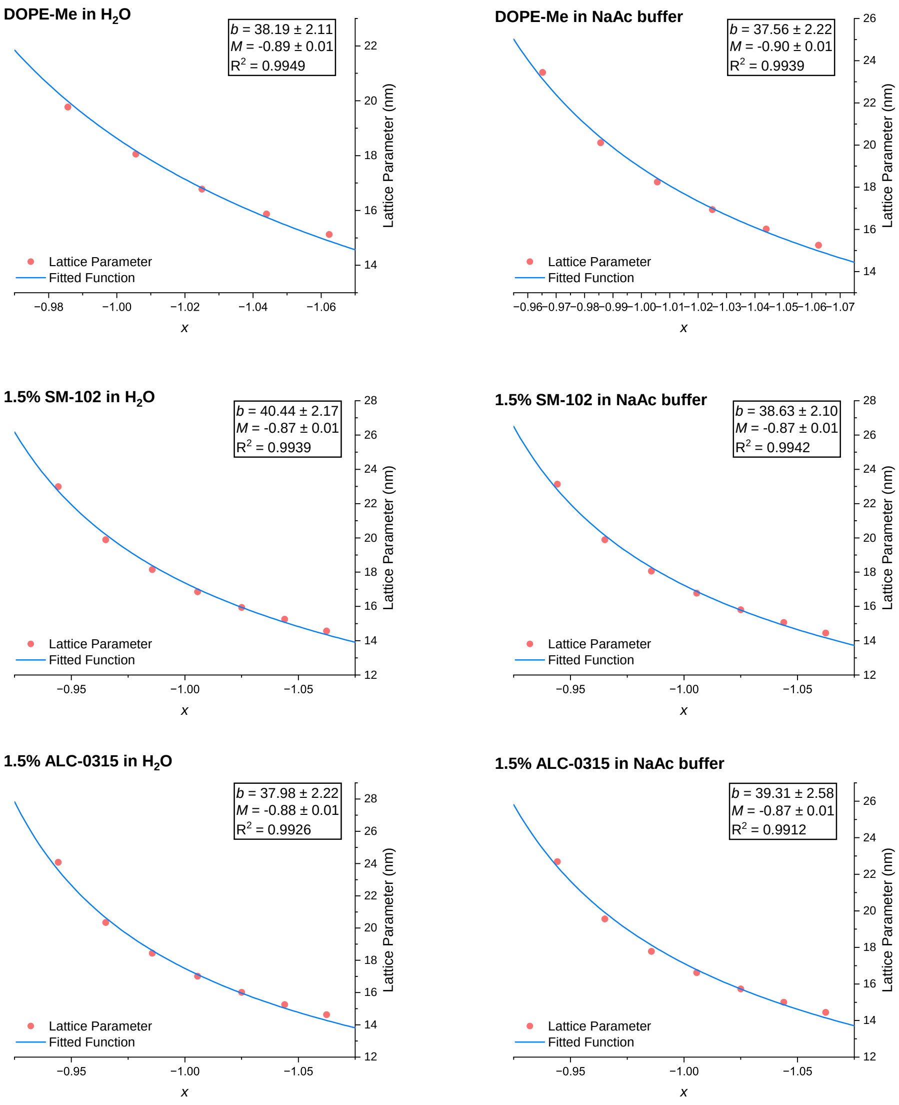


**Figure S12**: **Fitting for** $M$ **of DOPE-Me/IL Mixtures.** $M$ of DOPE-Me IL mixtures is fitted in the same way as Figure S8. The IL used are SM-102 and ALC-0315, which are lipids used in the COVID-19 vaccine formulation (*71*).

**Table S1**: **Sizes of the GMO/DOPC LNPs, early endosome membrane (EEM) vesicles and isolated endosomes.** Sizes measured by nanoparticle tracking analysis (NTA).

| Particle | Average Size (nm) |
|---|---|
| 0% GMO | 100.8 ± 1.8 nm |
| 15% GMO | 98.0 ± 3.5 nm |
| 35% GMO | 109.0 ± 0.8 nm |
| 65% GMO | 103.9 ± 2.2 nm |
| 85% GMO | 137.6 ± 0.2 nm |
| 100% GMO | 119.9 ± 3.2 nm |
| Isolated Endosomes | 137.2 ± 2.6 nm |
| EEM Vesicles | 103.0 ± 6.4 nm |

**Table S2**: The $\bar{\kappa}_m/\kappa_m$ $(M)$ values extracted and $Q$ calculated for DOPE-Me with small amounts of IL with different buffers at 37°C.

| DOPE-Me/IL Molar Ratio | Phase | Buffer | $M$ | $Q$ |
|---|---|---|---|---|
| 100/0 | $Im3m$ | $H_2O$ | −0.891 ± 0.007 | −0.049 ± 0.005 |
| 100/0 | $Im3m$ | NaAc buffer | −0.895 ± 0.007 | −0.054 ± 0.005 |
| 98.5/1.5 | $Im3m$ | $H_2O$ | −0.866 ± 0.007 | −0.024 ± 0.005 |
| 98.5/1.5 | $Im3m$ | NaAc buffer | −0.870 ± 0.007 | −0.028 ± 0.005 |
| 98.5/1.5 | $Im3m$ | $H_2O$ | −0.877 ± 0.007 | −0.035 ± 0.005 |
| 98.5/1.5 | $Im3m$ | NaAc buffer | −0.866 ± 0.009 | −0.024 ± 0.007 |
